# A high-throughput protein tagging toolkit that retains endogenous UTRs for studying gene regulation in Kinetoplastids

**DOI:** 10.1101/2024.11.02.621556

**Authors:** Carla Gilabert Carbajo, Xiaoyang Han, Bhairavi Savur, Arushi Upadhyaya, Fatima Taha, Michele Tinti, Richard J Wheeler, Phillip A. Yates, Calvin Tiengwe

## Abstract

Kinetoplastid parasites cause diseases that threaten human and animal health. To survive transitions between vertebrate hosts and insect vectors, these parasites rely on precise regulation of gene expression to adapt to environmental changes. Since gene regulation in Kinetoplastids is primarily post-transcriptional, developing efficient genetic tools for modifying genes at their endogenous loci while preserving regulatory mRNA elements is crucial for studying their complex biology.

We present a CRISPR/Cas9-based tagging system that preserves untranslated regulatory elements and uses a viral 2A peptide from *Thosea asigna* to generate two separate proteins from a single transcript: a drug-selectable marker and a tagged protein of interest. This dual-function design maintains native control elements, allowing discrimination between regulation of transcript abundance, translational efficiency, and post-translational events.

We validate the system by tagging six *Trypanosoma brucei* proteins and demonstrate: (i) high-efficiency positive selection and separation of drug-selectable marker and target protein, (ii) preservation of regulatory responses to environmental cues like heat shock and iron availability, and (iii) maintenance of stage-specific regulation during developmental transitions. This versatile toolkit is applicable to all kinetoplastids amenable to CRISPR/Cas9 editing, providing a powerful reverse genetic tool for studying post-transcriptional regulation and protein function in organisms where post-transcriptional control is dominant.

## Introduction

Kinetoplastid parasites, including *Trypanosoma brucei*, *T. cruzi*, and *Leishmania species*, cause significant vector-borne diseases in humans and animals. These parasites undergo complex developmental transitions as they switch between vertebrates and insect vectors, adapting to drastically different environments. Unlike most eukaryotes, kinetoplastids rely primarily on post-transcriptional gene regulation, with untranslated regions (UTRs) playing a crucial role in controlling mRNA fate and translation efficiency (1–3). Most protein-coding genes are transcribed by RNA Pol II in polycistronic units, with their expression controlled through mRNA stability and translation, as seen with heat shock protein 70 (Hsp70) regulated by 3′-UTR elements during the stress response (4–6), and the calpain-related proteins CAP5.5V and CAP5.5 during life cycle transitions (7, 8). Only a small subset of genes, including expression site-associated genes (ESAGs), undergo transcriptional regulation by RNA Pol I (9).

A prime example of UTR-mediated regulation is seen in ESAG6 and ESAG7, which encode subunits of the iron transporter, the transferrin receptor (TfR), in *T. brucei.* Their 3’ UTRs contain iron-responsive elements that enhance TfR mRNA stability in response to iron starvation (10, 11). Other important examples are procyclin and VSG, the major surface proteins of procyclic and bloodstream forms respectively, which are regulated by elements in their 3’ UTRs. Procyclin expression is controlled by several 3’ UTR elements, first identified by Hug et al (12), including a conserved 16-mer stem-loop structure (13) and additional regulatory sequences that affect RNA stability and translation (14–16). Conversely, the VSG mRNA 3’ UTR contains regulatory elements (17) including a 16-mer sequence essential for mRNA stability (18). More recently, a genome-wide analysis has predicted numerous UTR motifs and identified thousands of 3’ UTRs likely to influence mRNA abundance across different life cycle stages (19), highlighting the critical role of UTRs in enabling parasite adaptation.

Preserving UTRs is essential for studying gene function in kinetoplastids, as their disruption can alter expression patterns (10, 11). Several tagging methods have been developed for kinetoplastids, each with trade-offs between efficiency and preservation of regulatory elements. Early approaches, like long primer PCR (20, 21), were efficient, while others required cloning gene fragments for homologous recombination (22). PCR-only tagging in *T. brucei* and similar methods in *Leishmania* enabled scalable tagging but disrupt native UTRs when combined with drug-selectable markers (23). The TrypTag project, which tagged nearly 90% of the *T. brucei* proteome, has been a valuable resource for localisation studies, albeit with the recognised caveat that it replaces UTRs (24). CRISPR/Cas9 improves efficiency and precision but often introduces selection markers that interfere with endogenous regulation (25). Two CRISPR-based marker-free methods have been reported which preserve UTRs but require extensive subcloning by limiting dilution (26, 27). While these methods have advanced functional genomics, most still compromise regulatory elements, highlighting the need for tools that preserve UTR integrity while enabling efficient, high-throughput protein tagging and functional analysis.

First discovered in the Foot-and-Mouth Disease Virus (FMDV), the viral 2A peptide has become a powerful tool for expressing multiple proteins from a single open reading frame (ORF) (28, 29). Several versions of the 2A peptide have been identified across different viruses, each with similar functionality. These include FMDV (F2A), porcine teschovirus-1 (P2A), and *Thosea asigna* virus (T2A) (30). Interestingly, 2A-like sequences (L2A) have also been found in the L1Tc non-LTR retrotransposons of *T. cruzi*, *T. brucei*, *T. vivax* and *T. congolense,* which represents a rare example of these sequences occurring outside of a non-viral context (31). In general, the T2A peptide, with the sequence -EGRGSLLTCGDVEENPG↓P-, facilitates a mechanism known as “ribosomal skipping,” where the glycine-proline bond fails to form during translation, resulting in the production of two distinct proteins from the same transcript (28). This process, often called CHYSEL (cis-acting hydrolase element), “stop-and-go”, “self-processing”, or “co-translational cleavage”, will hereafter be referred to as “cleavage” for simplicity, with instances where the process fails being referred to as “uncleaved” proteins.

In Kinetoplastids, the T2A peptide has been shown to be highly efficient for co-expressing proteins under endogenous UTRs, enabling the study of post-transcriptional regulatory mechanisms. Some examples include the *L. mexicana* glucose transporter (LmxGT1) (32), the cortical cytoskeletal protein (KHARON1) (33) and the nucleotide transporter NT3 in *L. donovani* (34). The T2A system has also been applied for ectopic expression in *T. cruzi*, improving positive selection of transgenic parasites (35). These applications demonstrate the versatility of the T2A peptide system for studying protein localisation and regulatory mechanisms in kinetoplastids.

In this study, we combined CRISPR/Cas9 editing with the T2A peptide-based system to test its applicability in *T. brucei* by tagging two secretory proteins (ESAG3, ESAG7) which are under transcriptional and post-transcriptional regulation, and cytosolic Hsp70, part of a multi-gene array whose expression is regulated by its 3’ UTR in response to heat shock. We demonstrate efficient T2A peptide cleavage and regulation of tagged mRNA and/or proteins in their native loci. We also constructed new vectors with primer binding sites compatible with the plasmids for PCR-only tagging (pPOT series) (23) and designed an automated primer design tool to facilitate high-throughput tagging while preserving native UTRs. We validated the new vectors on three well-characterised proteins: glycosylphosphatidylinositol phospholipase C (GPI-PLC) (36) and two calpain-related proteins (CAP5.5V and CAP5.5) (7, 8), which show stage-specific expression, and confirmed that this differential expression was maintained in the tagged versions. Together, we provide a versatile tool that can be used at scale to study post-transcriptional and post-translational regulatory mechanisms in Kinetoplastids.

## Results

### Development of a CRISPR/Cas9-compatible T2A peptide tagging system for *T. brucei*

Our primary goal was to exploit the high efficiency of CRISPR/Cas9 genome editing to insert a “T2A peptide cassette” between the open reading frame (ORF) of a target gene and its native 5’ or 3’ UTR in *T. brucei*. The cassette contains a drug-selectable marker (drug^R^), a 2A peptide sequence (T2A), and a fluorescent protein or an epitope tag (Tag) (**Fig. 1**). We initially constructed donor plasmids in two configurations: drug^R^::T2A::Tag for insertion at the 5’ end (**Fig. 1A**), and Tag::T2A::drug^R^ for insertion at the 3’ end of the target gene (**Fig. 1B**). This design allowed for positive selection of transgenic parasites while preserving native mRNA processing signals within the UTRs, thus maintaining natural regulation. Available plasmids with fluorescent tags: mNeonGreen (mNG) or mScarlet (mSc), and drug-selectable markers: blasticidin (BSD), or puromycin (PAC) are shown in **Fig. S1**.

**Fig. 1.**
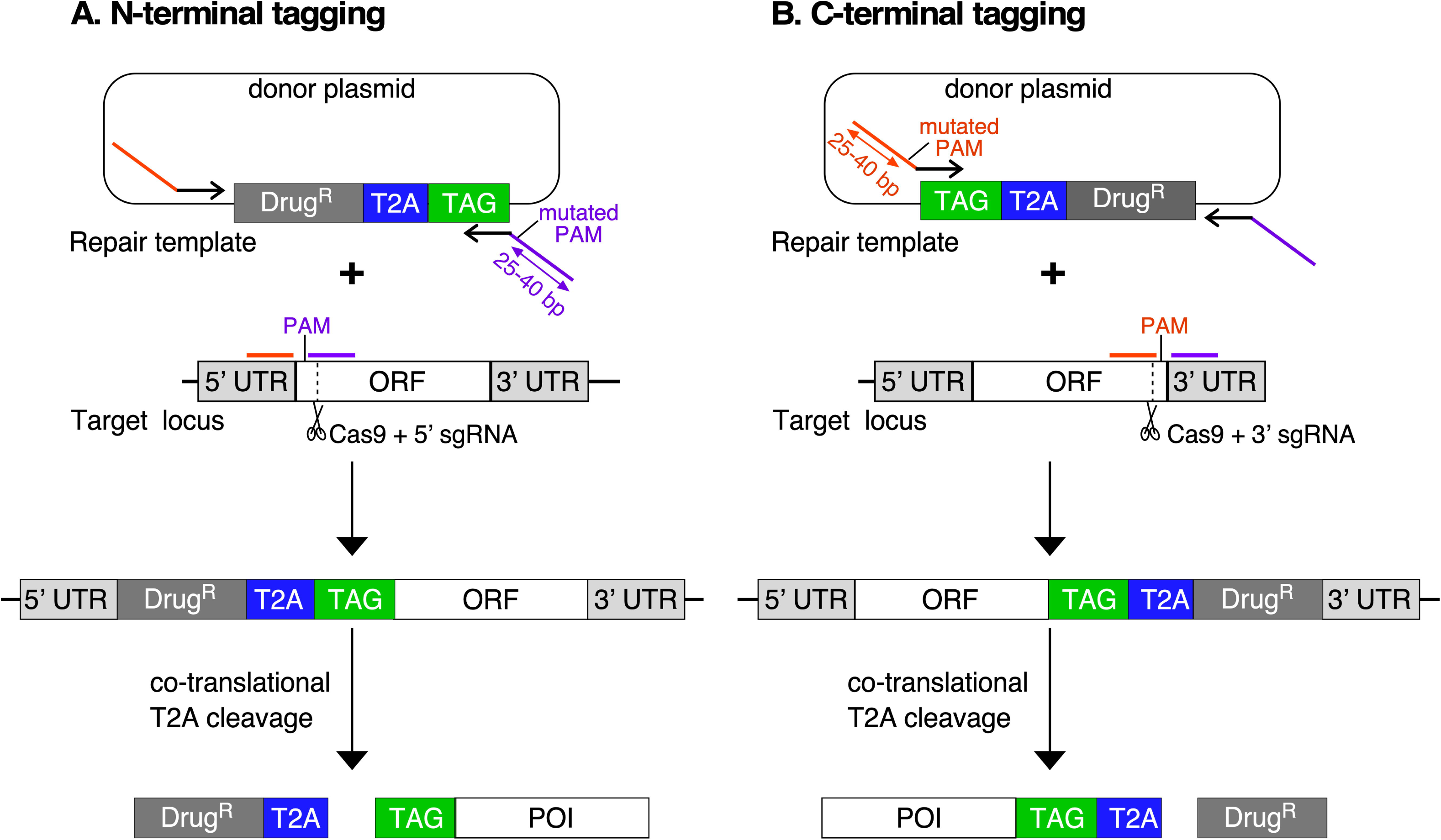
Strategy for endogenous T2A peptide-based protein tagging. The T2A peptide-based tagging system relies on CRISPR/Cas9-mediated insertion of a repair template either on the 5’ end (**A**) or the 3’end (**B**) of the open reading frame of a target gene. The repair template consists of a drug-selectable marker (Drug^R^) linked by a *Thosea asigna* T2A peptide sequence to a fluorescent protein or an epitope tag (TAG), placed at the N- or C-terminus of a protein of interest (POI). PCR amplification of donor repair templates uses primers containing ∼25 - 40 bp of homology arms (orange and purple lines) flanking the Drug^R^::T2A::TAG cassette. The single guide RNA (sgRNA) directs Cas9 to introduce a double-stranded break adjacent to the protospacer-adjacent motif (PAM), and mutated PAM sites from the repair template prevent further edits. Following correct integration, the T2A peptide causes co-translational cleavage, resulting in expression of both the Drug^R^ and POI in equal amounts. In both configurations, the sgRNA and repair template primers are designed so that the full-length endogenous 5’ and 3’ UTR sequences remain intact and drive expression of the Drug^R^ protein and POI.

For 5’-terminal tagging, the donor repair template drug^R^::T2A::Tag was amplified by PCR using primers containing 25-40 bp homology arms, targeting a protospacer adjacent motif (PAM) site near the start of the ORF. The ATG start codon was removed, but the 5’ UTR remained intact. For 3’-terminal tagging, the Tag::T2A::drug^R^ donor template was amplified using primers with homology arms to integrate the cassette before the stop codon, targeting a PAM site near the end of the ORF. This integration strategy incorporated 57 bp of plasmid backbone sequence between the targeting cassette and the native 3’ UTR, though the full-length 3’ UTR remained intact downstream of this insertion. In both cases the single-guide RNA (sgRNA) was generated by combining a gene-specific primer encoding a T7 promoter upstream, a 20-bp targeting sequence specifying the target site, and a 3’-end complementary to the sgRNA scaffold, as described in (37).

Both sgRNA and donor T2A cassettes were amplified by PCR using well-established protocols described by Beneke et al. (25). The PCR products were pooled for transfection into bloodstream form *T. brucei*, which constitutively express T7 RNA polymerase and Cas9. Cas9 introduces a double strand break at the target site, guided by the sgRNA designed near the PAM site at either the 5’ or 3’ end of the target gene. We incorporated mutated PAM sites into the repair template to prevent further Cas9 edits post-integration. Once integrated, the T2A peptide sequence mediates co-translational cleavage, allowing for independent expression of the drug resistance marker and the tagged protein of interest in equal amounts (38).

The donor plasmids described here were originally constructed for *Leishmania*, where we determined that T2A peptide conferred the highest cleavage efficiency among the various 2A peptides tested (**Fig S2**).

### Test case I: ESAG3 endogenous tagging shows efficient T2A cleavage and robust positive selection

To assess the efficiency of T2A peptide cleavage in *T. brucei* and demonstrate its utility for protein localisation studies, we targeted the *expression site associated gene 3* (*ESAG3*, gene IDs: Tb427.BES40.10, Tb427.BES40.16). *ESAG3* encodes an N-terminal putative ER targeting signal sequence with a predicted cleavage site between Ala22/Leu23 (SignalP v5.0). A two-component PCR amplicon mix was transfected into *T. brucei* T7/Cas9-expressing cells, consisting of a 20-bp sgRNA targeting adjacent to the PAM site near the stop codon and a donor mNG::T2A::PAC cassette with 40/30 bp homology arms. The stop codon was excluded from repair primers to allow in-frame fusion of mNG at the C-terminus, while synonymous mutations in the PAM site prevented further Cas9 edits.

Seven days post-transfection, genomic DNA PCR confirmed in-frame fusion of *ESAG3* ORF with mNG (expected size 594 bp), *PAC* with *ESAG3* 3’UTR (211 bp), *ESAG3* ORF with its native UTR (2.5 Kb) (**Fig 2A**). No PCR product was generated using genomic DNA from parental cells confirming specificity. To compare the relative mobility of the ESAG3::mT2A protein, we generated independent cell lines with C-terminally mNG-tagged ESAG3 fused with 6xTy (ESAG3::mTy) using the PCR-only tagging method (37) (**Fig 2B**). Western blot analysis revealed that the ESAG3::mT2A protein (69 kDa) migrated faster than ESAG3::mTy (76 kDa), indicative of efficient T2A cleavage. The expected MW of uncleaved ESAG3::mNG::T2A::PAC is 90 kDa. Since there are two copies of ESAG3 in the active expression site (BES1, Tb427_telo40_v2) in the Lister427 cell line (39), native anti-ESAG3 antibody immunoblots confirmed that tagging was single copy for most clones, as the untagged copy was detectable at the expected MW of ∼43 kDa (**Fig 2B)**. Flow cytometry showed that 93-95% of cells in the drug-selected clonal population were mNG-positive, confirming robust positive selection of correctly tagged cells (**Fig 2C**). Immunofluorescence microscopy showed a typical ER localisation pattern for ESAG3::mT2A, confirming proper targeting to the secretory pathway (**Fig 2D**).

**Fig. 2:**
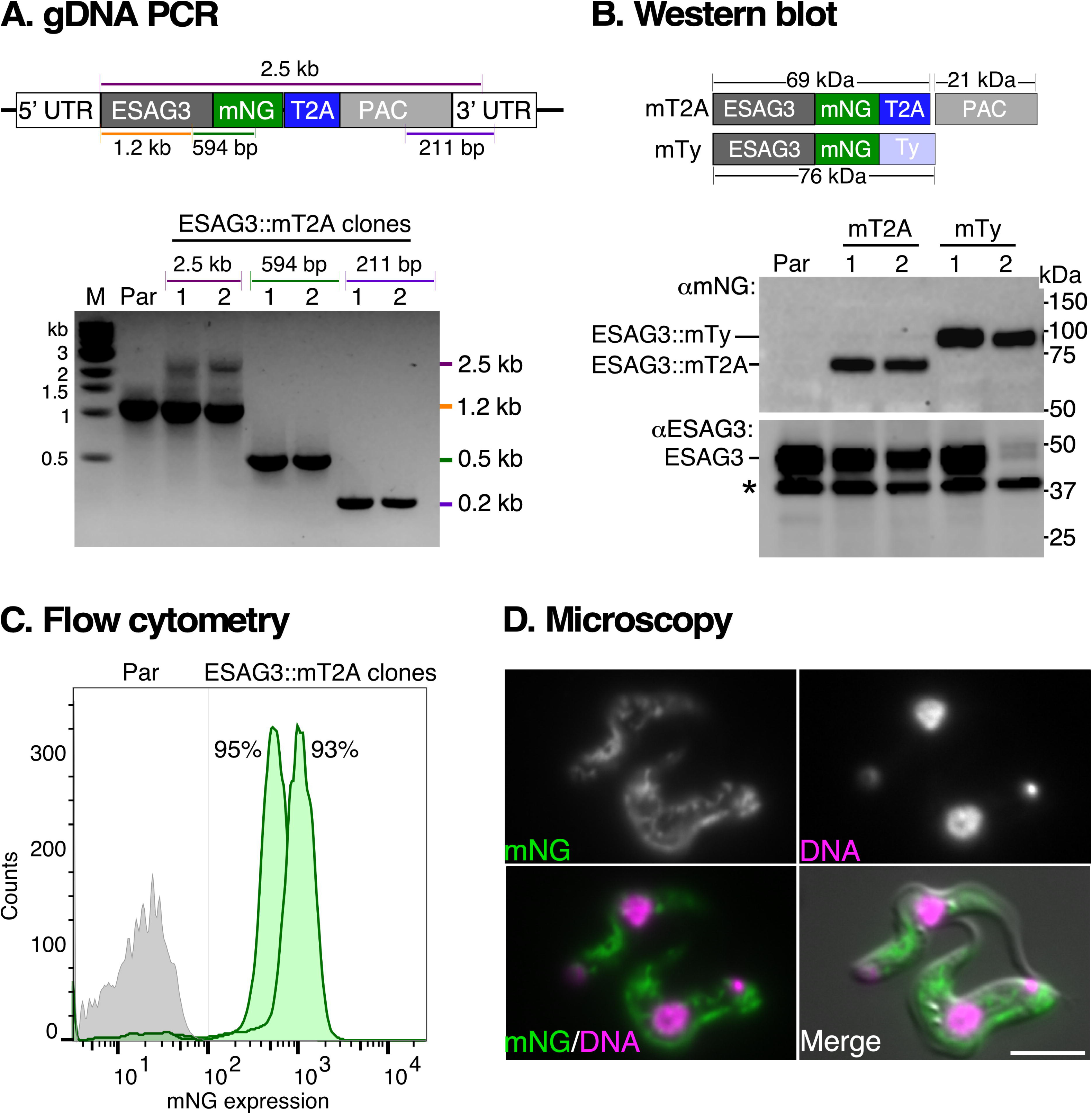
High-efficiency T2A cleavage in *T. brucei* and localisation of ESAG3::mNG::T2A. **A.** PCR validation of *ESAG3::mT2A* tagged genomic locus. Diagram shows the *ESAG3* gene locus after insertion of a repair cassette containing mNeonGreen (mNG), T2A peptide sequence (T2A), and puromycin (PAC) resistance gene for selection. PCR was performed using genomic DNA from parental cells (Par) and two transfected ESAG3::mT2A clones (1, 2). Coloured lines indicate positions of PCR primers to verify *ESAG3* locus-specific integration, with corresponding size fragments shown on the right of the agarose gel image. Molecular weight markers (M) are shown on the left. All diagrams are not drawn to scale. Note: The weaker intensity of the 2.5 Kb band reflects the expected reduced PCR efficiency for longer products. **B.** Validation of tagged ESAG3::mT2A protein. Top panel: Diagram shows predicted molecular weights (MW) without (90 kDa) and with (69 kDa) T2A cleavage during translation of *ESAG3::mT2A::PAC* (mT2A) mRNA. Independent cell lines were generated by CRISPR/Cas9 mediated fusion of ESAG3 with mNG with 6xTy (ESAG3::mTy, mTy, predicted MW 76 kDa) without incorporating a T2A peptide sequence. The ESAG3::mTy protein served as an internal control for SDS-PAGE migration analysis. Note that ESAG3::mTy was generated using the pPOT method which replaces the endogenous 3’ UTR with *PFR2* intergenic region. Bottom panel: Western blot analyses using anti-mNG (αmNG, top) and anti-ESAG3 peptide (αESAG3, bottom) antibodies. Asterisks indicate αESAG3 non-specific cross-reacting polypeptide. **C.** Evaluation of mNG-positive cells by flow cytometry. Parental cells (Par, gray histogram) and ESAG3::mNG::T2A-tagged cells (ESAG3::mT2A, clones 1 and 2, green histograms) are shown. Inset: percentage of mNG-positive cells. **D.** Localisation of ESAG3::mT2A protein visualised by live-cell epifluorescence microscopy (green), co-stained with Hoechst 33342 (magenta; DNA). Scale bar = 5 μM.

These results demonstrate precise successful insertion of the mNG::T2A::PAC cassette at the 3’ end of *ESAG3*, with highly efficient positive selection of tagged cells, and successful co-translational T2A cleavage in *T. brucei*. Furthermore, this tagging system enabled us to localise ESAG3 in bloodstream form trypanosomes, for the first time.

### Test case II: Endogenous T2A tagging of Hsp70 retains mRNA-level heat shock regulation

To further demonstrate the robustness of the T2A system, we targeted the cytosolic heat shock protein Hsp70. In *T. brucei*, Hsp70 is encoded by a tandem array of eight nearly identical genes (IDs: Tb427_110128800, Tb427_110128900 … Tb427_110129500) on chromosome 11 (40). In the Tb927 genome assembly, this array is collapsed into a single gene (Tb927.11.11330) (4, 41). Hsp70 mRNA stability is regulated by heat response elements in the 3′-UTR during heat shock (4–6), but corresponding changes at the protein level have not been demonstrated. Our goals were to investigate the behaviour of endogenously T2A-tagged Hsp70 following heat shock and to assess whether our system facilitated single-copy integration within a multi-gene array.

We designed primers based on the Tb927 genome copy to tag Hsp70 at both the N- and C-termini with mScarlet (mSc) and mNeonGreen (mNG), respectively, under puromycin selection (**Fig. 3A**). The design of sgRNA and donor repair templates followed the strategy used for ESAG3. A GSGS linker was added between the mSc tag and the *Hsp70* ORF for N-terminal tagging, and 25 bp homology arms were used in both repair templates. Post-transfection, locus-specific PCR genotyping of two clones produced products of sizes consistent with the predicted fusions: *Hsp70* ORF with mNG (409 bp), *PAC* with the *Hsp70* 3’ UTR (304 bp), and *Hsp70* 5’ UTR with *PAC* (339 bp) (**Fig. 3A**). No products were generated from genomic DNA purified from parental cells, confirming locus-specific modification.

**Fig. 3:**
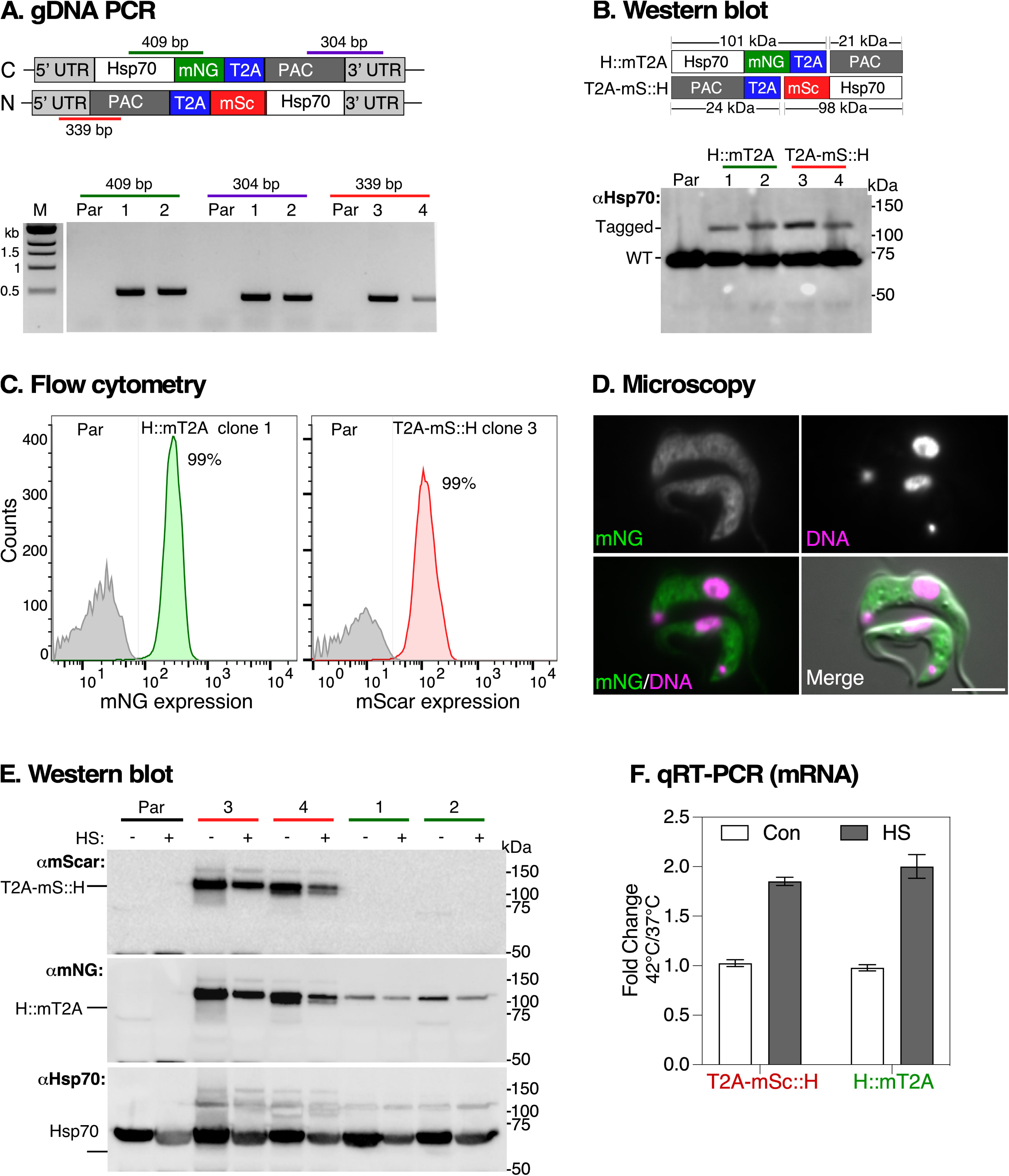
Effect of heat shock on T2A-tagged Hsp70 mRNA and protein following heat shock. **A.** PCR validation of *Hsp70* tagged loci. Top panel shows schematic of the locus after insertion of a mNG::2A::drug repair cassette. Bottom panel shows agarose gel image of PCR products using genomic DNA extracted from parental (Par) cells or clonal cells transfected with repair templates for Hsp70 tagged either at the C-terminus with mNeonGreen (mNG, Hsp70::mT2A) or at the N-terminus with mScarlet (mSc, T2A-mS::Hsp70). Coloured lines indicate positions of PCR primers with corresponding size fragments shown above each lane on the agarose gel image. M: molecular markers, two clones for mNG (1, 2) and mSc (3, 4). **B.** Top panel: Diagram shows the predicted MWs of tagged Hsp70 protein before (122 kDa) and after (101 kDa) T2A-mediated co-translational intra-ribosome skipping. Bottom panel: Western blot analyses of tagged and untagged (WT) Hsp70 protein with native anti-Hsp70 (αHsp70) antibody. **C.** Flow cytometry analysis of tagged Hsp70 cells. Parental cells (Par, gray) and cells in which Hsp70 was either mNG-tagged (left) or mSc-tagged (right). Inset shows percentage of positive fluorescent cells. **D.** Localisation of Hsp70::mT2A protein visualised by live-cell epifluorescence microscopy (green), co-stained with Hoechst 33342 (magenta; DNA). Scale bar = 5 μM. **E.** Regulation of tagged Hsp70 protein expression after heat shock (HS). Cell cultures were maintained at 37 °C (-) or transferred to 42 °C (+) for one hour. Total protein extracts were analysed by western blot with anti-mSc (αmScar), anti-mNG (αmNG) and anti-Hsp70 (αHsp70) antibodies. Two clones are shown for parental (Par) and each tagged cell line. Note: The same membrane was sequentially probed without stripping, starting with αmScar, followed by αmNG, and finally αHsp70. **F.** Regulation of *Hsp70* mRNA expression after heat shock. Cell cultures were maintained at 37 °C (Control, Con) or transferred to 42 °C (Heat shock, HS) for one hour and mRNA levels quantified by qRT-PCR using primers specific to endogenous Hsp70. Fold change is calculated relative to 37 °C controls with *ZFP3* as endogenous control. Error bars represent standard deviation from two clonal cell populations with three technical duplicates for each clone.

Western blot analysis with anti-Hsp70 antibodies showed two bands in all selected clones: one at ∼70 kDa (untagged endogenous Hsp70) and another at ∼100 kDa, (tagged Hsp70: 101 kDa for Hsp70::mT2A and 98 kDa for T2A-mSc::H). The higher MW bands were absent in parental cells (**Fig. 3B**), indicating successful tagging. Flow cytometry revealed that ∼99% of cells were positive for either mNG or mSc (**Fig 3C**), and live-cell microscopy confirmed the characteristic cytosolic localisation of Hsp70::mT2A (**Fig. 3D**), indicating that T2A tagging did not cause Hsp70 mis-localisation.

To assess whether expression of mT2A-tagged Hsp70 tracked with the untagged allele following heat shock, we performed immunoblotting with anti-mScarlet, anti-mNG, and anti-Hsp70 antibodies. We observed a slight reduction in both tagged and endogenous Hsp70 protein levels after heat shock (**Fig. 3E**). However, qRT-PCR analysis showed a 2-fold upregulation of Hsp70 mRNA (**Fig. 3F**), suggesting a typical heat shock response at the mRNA level (6). We attribute the reduction in protein levels to either global translational arrest or cell stress resulting from the heat shock treatment (42).

Overall, our results confirmed UTR-dependent *Hsp70* mRNA regulation after heat shock (4), but not at the protein level - a pattern consistent with established models in *T. brucei* where selective mRNA stabilisation enables continued Hsp70 synthesis during stress-induced translational repression (5, 6). We successfully achieved efficient single-copy tagging of cytosolic Hsp70 at both the N- and C-termini within a complex multi-gene array. The apparent lower expression of tagged Hsp70 compared to the wild type likely indicates that only one of the eight nearly identical gene copies was tagged, accounting for approximately 1/16th of the total signal in Western blots of a diploid genome. We confirmed this using whole genome sequencing analysis and found no evidence of deletions of any Hsp70 genes in the array (**Fig. S3**).

### Test case III: T2A-tagged ESAG7 preserves iron-dependent regulation and function

In functional studies, it is essential that tagging does not disrupt protein function or its regulatory elements at the RNA level. In *T. brucei,* ESAG7 (E7) exemplifies this dual requirement: at the protein level, it forms a 1:1 stoichiometric complex with ESAG6 to create a functional transferrin receptor (TfR) for iron uptake (43). At the RNA level, its expression is regulated by iron-responsive elements in its 3’ UTR that control mRNA stability in response to iron availability (44). Tagging approaches that replace the native 3’ UTR can disrupt this iron-dependent regulation, potentially altering the expression levels and stoichiometry of ESAG6 and ESAG7, thereby interfering with TfR function (10, 11).

To investigate regulation and function of ESAG7, we generated mNG::T2A- and mNG::6xTy-tagged ESAG7 cell lines (E7::mT2A and E7::mTy, respectively) (**Fig 4A**). E7::mT2A tagging retained the native *E7* 3’ UTR, while the E7::mTy tagging replaced the 3’ UTR with the *PFR2* intergenic region. Hsp70 (H::mT2A) was also tagged as a control. All analyses were performed after treatment with the iron chelator Deferoxamine (DFO), which induces iron starvation. qRT-PCR analysis showed 2-fold increased mNG transcript levels in E7::mT2A cells following DFO treatment, while E7::mTy showed a reduction, and H::mT2A cells remained unchanged (**Fig 4B**). This pattern was reflected at the protein level, as Western blotting confirmed increased E7::mT2A protein abundance, with E7::mTy and H::mT2A remaining unaffected (**Fig 4C**). Flow cytometry further supported these findings, showing a 3-fold increase in mNG expression in E7::mT2A cells, while no significant changes were observed in E7::mTy or H::mT2A (**Fig 4D**).

**Fig. 4:**
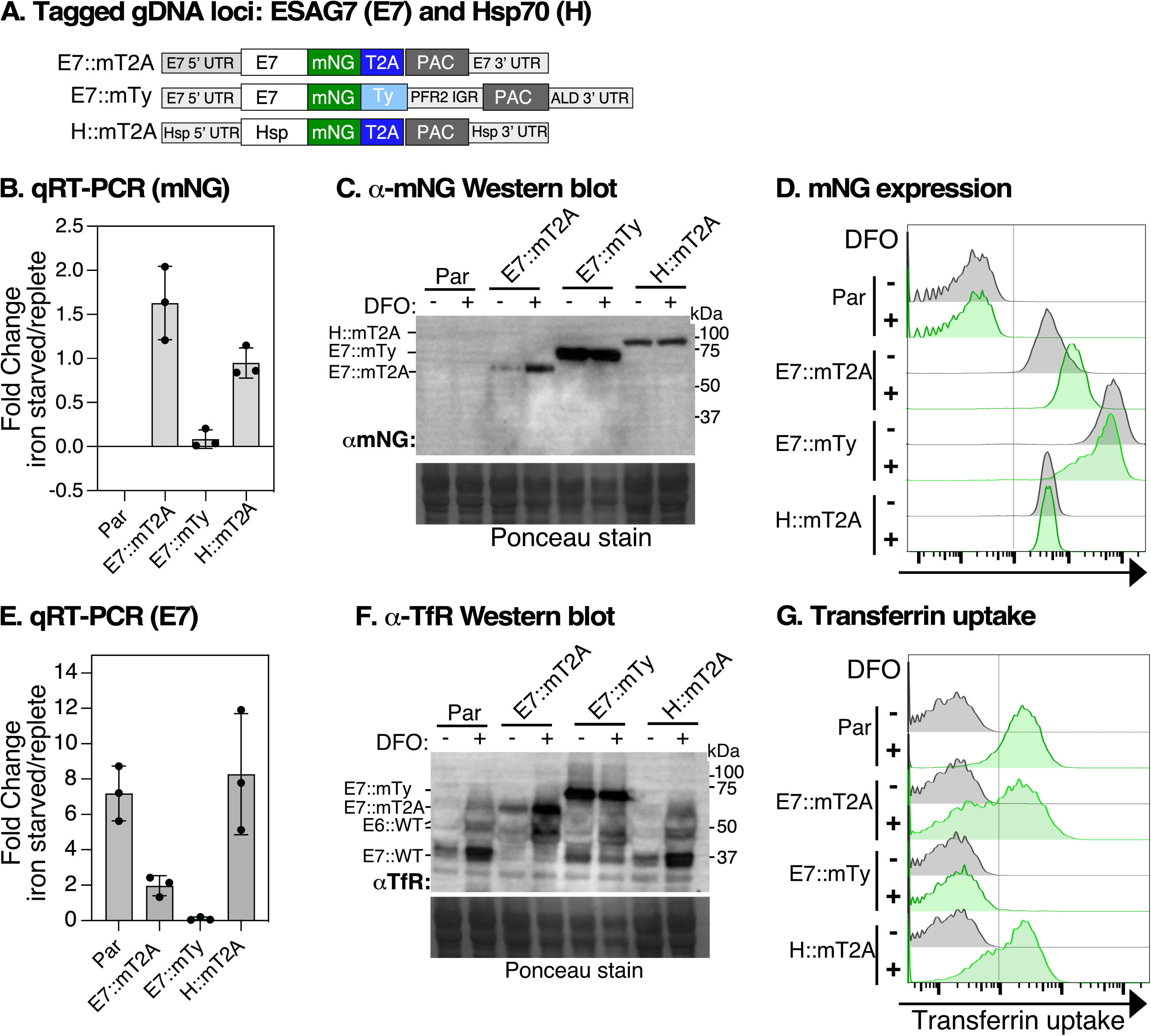
Endogenous T2A tagging of ESAG7 maintains iron-dependent regulation. **A.** Schematic shows the modified *ESAG7* (*E7)* and *Hsp70* genomic DNA locus after insertion of the mNG::2A::PAC repair cassette between the ORF and their native 3’ UTRs (E7::mT2A and H::mT2A) or insertion of mNG::6xTy with *paraflagellar rod protein 2* intergenic region *(PFR2* IGR; E7::mTy). E7::mT2A and E7::mTy tagging preserve endogenous 5’ UTRs. Diagrams not drawn to scale. **B.** qRT-PCR analysis of *mNG* transcript abundance in cells expressing E7::mT2A, E7::mTy, or H::mT2A following treatment with the iron chelator Deferoxamine (DFO). Transcript levels were normalised to *ZFP3* signal and expressed as a ratio to control treatment (iron-replete) values. Data are presented as means ± SD, n = 3 independent experiments from a single clone, with 2 technical replicates for each n. **C.** Western blot analysis of E7::mT2A protein levels using a-mNG antibody in different cell lines under iron-replete (DFO, -) and iron-starved (DFO, +) conditions. Ponceau staining indicates protein loading. **D.** Flow cytometry analysis of mNG expression in different cell lines under iron-replete (DFO, -) and iron-starved (DFO, +) conditions. Histograms show fluorescence intensity indicating mNG expression levels. **E.** qRT-PCR analysis of *E7* transcript levels in parental (Par) cells or cells expressing E7::mT2A, E7::mTy, or H::mT2A in iron-starved (DFO, +) conditions expressed as a ratio to control treatment (iron-replete) values and normalised to *ZFP3* signal. **F.** Western blot analysis of total protein using α-TfR antibody to detect total TfR expression in whole cell extracts. TfR is a heterodimer of ESAG6 (E6) and ESAG7 (E7). In parental (Par) and H::mT2A lanes, the predicted MW of endogenous E7 (E7::WT) is 37 kDa, while E6 (E6::WT) migrates as two species: ∼44 kDa (un-glycosylated) and >50 kDa (fully glycosylated) – see lanes 1 and 2 (par) and Lanes 7 and 8 (H::mT2A). The predicted MW of “cleaved” E7::mT2A, E7::mTy, H::mT2A are 67 kDa, 74 kDa, and 110 kDa respectively (lanes 3, 4, 5, and 6). Par and H::mT2A tagged cells serve as a control for DFO treatment. Ponceau-stained membrane shows loading control. The blot is representative of n = 3 independent experiments from one clone. **G.** Flow cytometry analysis of transferrin uptake in different cell lines under iron-replete (DFO, -) and iron-starved (DFO, +) conditions. Histograms show fluorescence intensity indicating transferrin uptake levels.

The functional implications of these findings were assessed by analysing native TfR levels and transferrin (Tf) uptake. Both qRT-PCR and Western blot analyses revealed increased TfR levels in parental, H::mT2A, and E7::mT2A cells after DFO treatment, but showed a significant reduction in E7::mTy cells (**Fig 4E** and **4F**). Tf uptake assays demonstrated significant increased uptake in E7::mT2A cells, confirming functional TfR activity, whereas E7::mTy cells showed no Tf uptake, indicating a disruption of iron-dependent regulation and receptor function (**Fig 4G**).

Collectively, these data demonstrate that mNG::T2A-tagged ESAG7, when retaining its native 3’ UTR, maintains iron-dependent regulation and function. In contrast, replacing the 3’ UTR with the *PFR2* intergenic region disrupts both iron-responsive regulation and TfR function, likely by perturbing the coordinated expression of ESAG7 with its heterodimeric partner ESAG6 required for functional receptor assembly. This validates our T2A-tagging approach as an effective tool for studying post-transcriptional iron regulation and hence likely other regulatory systems in *T. brucei*.

### Confirmation of site-specific integration and in-frame fusion of the T2A gene tagging by whole genome sequencing

Finally, to confirm site-specific integration and in-frame fusion of mNG::T2A with the target ORFs, we performed paired-end whole genome sequencing (WGS) of the five tagged cell lines at 30X genome coverage. The reads were mapped to either the wild-type genomic locus or a manually reconstructed locus containing the mNG::T2A or mNG::6xTy tagging cassettes fused to the target genes (**Fig. S4**). In all cases, WGS confirmed that the donor repair cassettes were inserted in-frame with the target gene in tagged cells, with no frameshifts or premature stop codons observed, confirming site-specific in-frame fusion. Collectively, combining data from WGS, integration PCRs, and Western blot analyses, we conclude that mNG::T2A fused to target ORFs predominantly as single-copy integrations, and that T2A peptide-mediated cleavage occurs with high efficiency. Collectively these results show that this approach is highly suitable for epitope-tagging secretory and cytosolic proteins at their endogenous loci in *T. brucei*.

### Dedicated modular plasmid templates for high-throughput CRISPR/Cas9 T2A tagging: The pRExT2A system

To enhance the efficiency and versatility of T2A tagging in high-throughput applications, we developed the pRExT2A system - plasmids for *R*egulated co-*Ex*pression mediated by *T2A* peptide) - a set of dedicated plasmid templates optimised for *T. brucei*. This new design addresses two key limitations of the initial T2A tagging system where: (i) the coding sequences for *mNG*, *mSc*, and *drug^R^* were optimised for *Leishmania* codon usage, which differs from *T. brucei* and could lead to inefficient translation and reduced expression levels; and (ii) the addition of 57 bp to the 3’ UTR during C-terminal tagging, which could potentially disrupt native mRNA processing signals and affect gene regulation.

To overcome these issues, we designed pRExT2A to be fully compatible with the pPOT (plasmids for PCR-only tagging) platform by incorporating the coding sequences for drug resistance markers and fluorescent proteins from pPOT into our new plasmids (23). Our goal was to integrate the efficiency, modularity, and cloning-free advantages of pPOT while ensuring that primer usage is compatible across both systems.

### Features and primer design of the pRExT2A system

The pRExT2A system features modular N- and C-terminal tagging donor plasmids encoding either a 3xTy-tagged fluorescent protein or a 3xTy epitope tag, linked via the T2A peptide to a 3xTy-tagged drug^R^ (**Fig. 5**). For N-terminal tagging, the constructs follow the format: drug^R^::3xTy::T2A::3xTy::FP or drug^R^::3xTy::T2A::3xTy (**Fig. 5A**), while for C-terminal tagging the orientation was as follows: FP::3xTy::T2A::drugR::3xTy or 3xTy::T2A::drug^R^::3xTy (**Fig. 5B**).

**Fig. 5.**
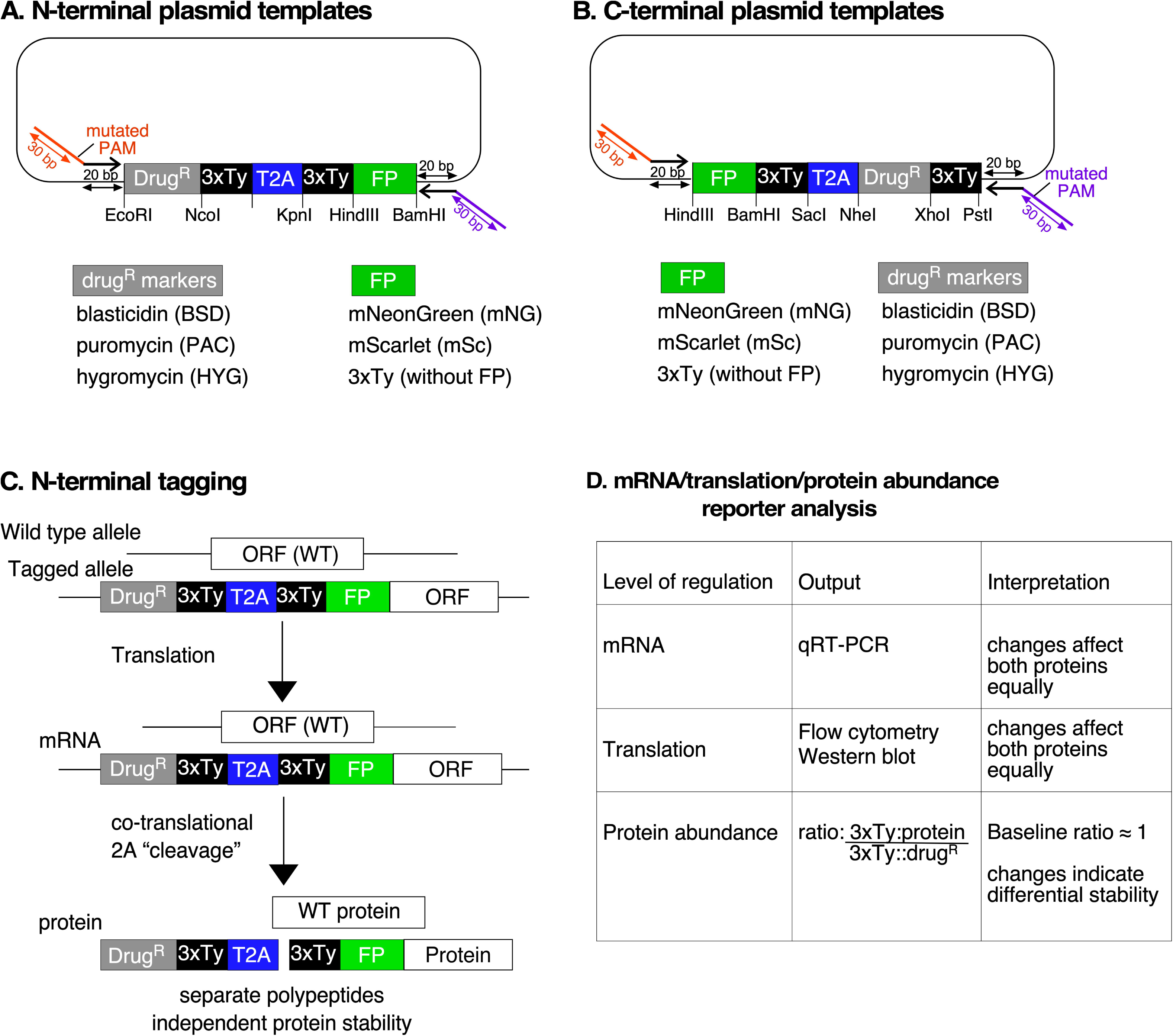
Map of pRExT2A modular template plasmids and strategy for studying post-transcriptional and post translational gene regulation. The schematic illustrates the design of CRISPR/Cas9-mediated modular donor vectors for N-terminal (**A**) and C-terminal (**B**) T2A tagging of target genes in *T. brucei*. The repair cassette includes a drug resistance marker (drug^R^, BSD, PAC, HYG), a 3xTy epitope, a T2A peptide, and a fluorescent protein tag (FP), such as mNeonGreen (mNG) or mScarlet (mSc). 3xTy epitope tags, drug^R^, and FP are flanked by unique restriction sites (EcoRI, NcoI, HindIII, etc.) to facilitate cloning and customisation. Arrows indicate the positions of forward and reverse primers used for the amplification of repair templates. The forward and reverse primers contain a 20 bp binding site to donor plasmid and 30 bp homology arms matching the target sequence containing a mutated PAM site, as described in Fig.1. The plasmid backbone is pUC57-simple (GenScript), and the full list of sequences is deposited in Addgene. **C.** The process of N-terminal tagging is depicted, where CRISPR/Cas9-mediated insertion of the repair cassette replaces one wild-type (WT) allele with a tagged allele at the 5’ end of the target gene. During translation, the T2A sequence facilitates co-translational cleavage, producing two distinct polypeptides: one containing the 3xTy-tagged drug^R^ marker and another containing the 3xTy-tagged FP-tagged protein. This strategy ensures that both the drug^R^ and FP-tagged proteins are independently stable, allowing for accurate analysis of gene regulatory mechanisms. **D.** The table outlines the different levels of regulation that can be studied using this system. The mRNA level can be analysed using qRT-PCR. At the translation level, protein expression can be monitored using flow cytometry or Western blot to detect 3xTy-tagged FP-tagged protein and the 3xTy-tagged drug^R^ marker. By measuring relative protein abundance of 3xTy-tagged protein to the 3x-tagged drug^R^ marker before and after stimulus exposure using the drugR as an internal reference, one can distinguish between regulation at the transcript level (where both proteins change similarly) and differential protein stability (where the target protein changes independently).

The primer design workflow remains consistent with our previous protocol (**Fig. 1**), but we modified the primer binding sites to match the pPOT system. This allows users to use identical primer pairs and sgRNAs across both systems. Although an improvement over the original T2A tagging cassettes, this design introduces a short 20 bp scar into the 5’ or 3’ UTR after integration - depending on whether N- or C-terminal tagging is used (**Fig. 5A** and **5B**). To facilitate high-throughput applications, we developed an automated primer design tool [https://github.com/zephyris/scarlesstagging] which generates primers compatible with both pRExT2A and pPOT systems. This tool significantly simplifies experimental setup and minimises design errors. To facilitate adoption, we provide pre-designed primers (including scarless options) for all *T. brucei* genes in TREU927, Lister 427 strains, and selected species, available at: https://github.com/zephyris/scarlesstagging/tree/main/examples/py/tritrypdb-genomes-v68.

### Dual functionality of the 3xTy-tagged drug^R^

A key feature of the pRExT2A system is the inclusion of a 3xTy-tagged drug^R^ which serves two functions: as a selection marker controlled by the endogenous UTRs of the target gene and an internal reference for measuring relative protein abundance before and after stimulus exposure (**Fig. 5C)**. Since the drug^R^ and the target protein are translated from the same mRNA but yield distinct proteins, changes in mRNA abundance or translational efficiency affect both proteins proportionally. Comparing protein levels using Western blotting or flow cytometry reveals whether regulation occurs at the transcript level (where both proteins change similarly) or through differential protein stability (where the target protein changes independently of the drug^R^) (**Fig. 5C** and **5D**). This design enables researchers to determine the level of regulation by measuring mRNA abundance (using qRT-PCR) and protein levels within the framework of a single experiment. A constant protein/drug^R^ indicates regulation at the mRNA or translational level, while changes in ratios from baseline reflect differential protein stability (**Fig 5D**), as exemplified by T2A-based reporter systems in *Leishmania* (32, 34).

### Modularity of the pRExT2A System

We developed plasmid versions with various drug-selectable markers, including blasticidin (BSD), hygromycin (HYG), and puromycin (PAC), as well as two fluorescent proteins, mNeonGreen (mNG) and mScarlet (mSc) (**Table 1**). This modular design allows for easy swapping of tags and drug^R^ markers, providing flexibility to accommodate different experimental needs.

**Table 1:** pRExT2A tagging plasmid templates for N- and C-terminal tagging.

| Name | Tag | DrugR | Configurations |
| --- | --- | --- | --- |
| **N-terminal tagging templates** |  |  |  |
| pRExT2A-NT-BmNG | 3xTy::mNG | BSD::3xTy | BSD::3xTy::T2A::3xTy::mNG |
| pRExT2A-NT-BmSc | 3xTy::mSc | BSD::3xTy | BSD::3xTy::T2A::3xTy::mSc |
| pRExT2A-NT-BTy | 3xTy | BSD::3xTy | BSD::3xTy::T2A::3xTy |
| pRExT2A-NT-PmSc | 3xTy::mSc | PAC::3xTy | PAC::3xTy::T2A::3xTy::mSc |
| pRExT2A-NT-PmNG | 3xTy::mNG | PAC::3xTy | PAC::3xTy::T2A::3xTy::mNG |
| pRExT2A-NT-PTy | 3xTy | PAC::3xTy | PAC::3xTy::T2A::3xTy |
| pRExT2A-NT-HmNG | 3xTy::mNG | HYG::3xTy | HYG::3xTy::T2A::3xTy::mNG |
| pRExT2A-NT-HTy | 3xTy | HYG::3xTy | HYG::3xTy::T2A::3xTy |
| **C-terminal tagging templates** |  |  |  |
| pRExT2A-CT-BmNG | mNG::3xTy | BSD::3xTy | mNG::3xTy::T2A::BSD::3xTy |
| pRExT2A-CT-BmSc | mSc::3xTy | BSD::3xTy | mSc::3xTy::T2A::BSD::3xTy |
| pRExT2A-CT-BTy | 3xTy | BSD::3xTy | 3xTy::T2A::BSD::3xTy |
| pRExT2A-CT-PmSc | mSc::3xTy | PAC::3xTy | mSc::3xTy::T2A::PAC::3xTy |
| pRExT2A-CT-PmNG | mNG::3xTy | PAC::3xTy | mNG::3xTy::T2A::PAC::3xTy |
| pRExT2A-CT-PTy | 3xTy | PAC::3xTy | 3xTy::T2A::PAC::3xTy |
| pRExT2A-CT-BmSc | mSc::3xTy | HYG::3xTy | mSc::3xTy::T2A::HYG::3xTy |
| pRExT2A-CT-BTy | 3xTy | HYG::3xTy | 3xTy::T2A::HYG::3xTy |

### The pRExT2A system retains life cycle stage-specific gene regulation and confirms UTR-mediated control

To evaluate the ability of the pRExT2A system to preserve critical regulatory elements, we tested three developmentally regulated genes in *T. brucei*: glycosylphosphatidylinositol phospholipase C (GPI-PLC), which is expressed exclusively in bloodstream forms via 3’ UTR regulatory elements, and two paralogous calpain-related proteins, CAP5.5V (bloodstream-specific) and CAP5.5 (procyclic-specific) (7, 8, 45). These genes provide ideal test cases for assessing the system’s ability to maintain post-transcriptional regulatory mechanisms during *in vitro* differentiation.

#### GPI-PLC example

GPI-PLC (Tb927.2.6000) is expressed exclusively in bloodstream forms via regulatory elements in its 3’ UTR, and it’s enzymatic activity is essential for VSG shedding during the early stages of differentiation to procyclic forms (45, 46). To track its expression during bloodstream to procyclic form differentiation, we generated C-terminal mNG::3xTy fusions (GPI-PLC::mT2A::3xTy) using the pRExT2A system, under hygromycin selection (**Fig. 6A)**. Live imaging showed GPI-PLC::mT2A::3xTy predominantly localised to the flagella membrane (**Fig. 6B**), consistent with native GPI-PLC antibody staining (47) and GPI-PLC-eYFP localisation (48). This pattern differs from the dual flagella/cell body distribution seen with pPOT tagging where the 3’-UTR was replaced (25). Western blot analysis using validated anti-GPI-PLC antibodies (47), showed expression of both native GPI-PLC (∼41 kDa) and GPI-PLC::mT2A::3xTy (G::mT2A; ∼73 kDa) specifically in bloodstream form cells (**Fig 6C, left**). The stoichiometric ratio between tagged and wild-type alleles indicated single-copy integration in the diploid genome, and efficient T2A cleavage was confirmed by detection of separate 3xTy::HYG product (42 kDa) in anti-Ty Western blots (**Fig 6C, right**). Although the tagged and untagged alleles in the G::mT2A cell line showed lower signal intensity with the anti-GPI-PLC antibody compared to the parental cell line (22, 48), the tagged protein maintained correct developmental regulation and localisation. Importantly, flow cytometry showed that GPI-PLC::mT2A expression dropped to near-background levels after differentiation to procyclic forms (**Fig 6D, 6E**), indicating that the pRExT2A system preserves the native UTR-mediated regulation of GPI-PLC expression. The distinct localisation patterns between our UTR-preserved and UTR-replaced constructs (25) highlight the importance of maintaining endogenous UTRs for proper protein expression.

**Fig. 6:**
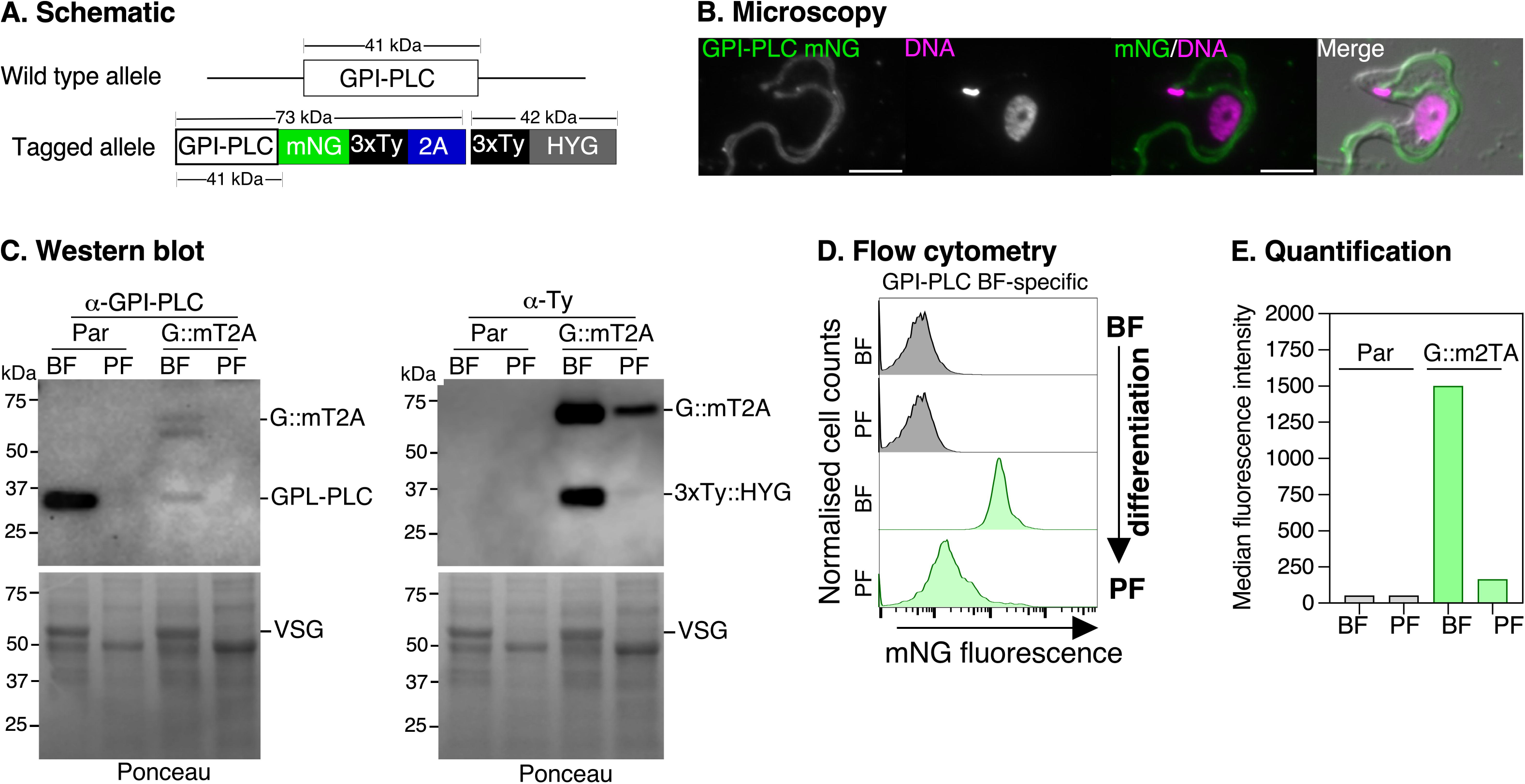
Tagging of GPI-PLC preserves life cycle-dependent expression patterns during bloodstream (BF) to procyclic form (PF) differentiation. **A.** Schematic of the GPI-PLC protein tagged with mNG::3xTy, fused via the T2A peptide to a Hygromycin resistance protein (3xTy::HYG) at its C-terminus. The approximate MW (in kDa) of the tagged and untagged proteins are indicated above the schematics. **B.** Localisation of GPI-PLC::mT2A protein visualised by live-cell epifluorescence microscopy (green), co-stained with Hoechst 33342 (magenta; DNA). Scale bar = 5 μM. **C.** Western blot analysis of GPI-PLC expression in parental (Par) and GPI-PLC::mT2A-tagged cells in BF and PF cells. The α-GPI-PLC antibody confirms BF-specific GPI-PLC expression showing stoichiometric expression of endogenous (GPI-PLC) and tagged GPI-PLC (G:mT2A). Ponceau staining showing a band corresponding to the MW of variant surface glycoprotein (VSG) in bloodstream but not in procyclic forms. α-Ty antibody (right) detected GPI-PLC::mT2A (G::mT2A) and 3xTy::HYG (HYG) proteins in BF cells, with signals at the expected size (∼73 kDa). No signal was observed in PF cells. **D.** Flow cytometry analysis of mNG fluorescence in GPI-PLC::mT2A-tagged cells during *in vitro* differentiation of BF to PF stages. BF cells display high fluorescence, confirming GPI-PLC expression, whereas PF cells show minimal fluorescence. Parental cells (Par) serve as a negative control. **E.** Quantification of flow cytometry data in D, showing median fluorescence intensity for mNG in BF and PF stages.

#### CAP5.5V and CAP5.5 examples

CAP5.5V (Tb927.8.8330) and CAP5.5 (Tb927.4.3950) play crucial roles in maintaining cell morphology by organising the subpellicular cytoskeleton in bloodstream and procyclic forms, respectively (8). In the TrypTag project, CAP5.5V showed no expression in procyclic forms when tagged at the N-terminus, but C-terminal tagging, which replaced the native 3’ UTR, led to inappropriate expression in procyclic forms, demonstrating the critical role of UTR-mediated regulation (49). To prevent such dysregulation, we preserved the native UTRs while tagging CAP5.5V and CAP5.5 at their C-termini using the pRExT2A system (**Fig. 7A)**.

**Fig. 7:**
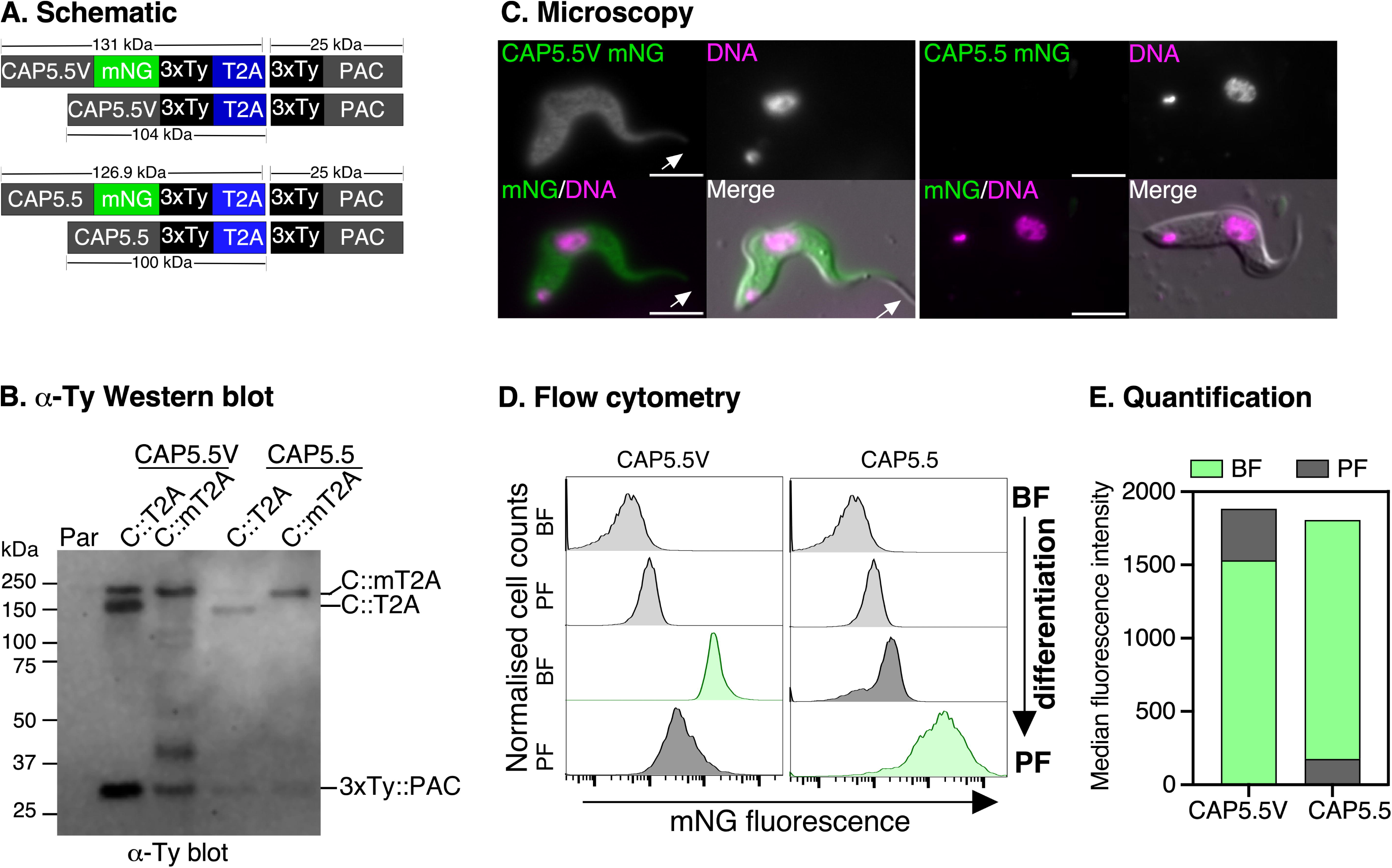
Tagging of CAP5.5V and CAP5.5 confirms stage-specific expression during bloodstream to procyclic form differentiation. **A.** Schematic of CAP5.5V and CAP5.5 proteins tagged with mNG::3xTy::2A::3xTy::PAC or 3xTy::2A::3xTy::PAC at their C-termini. The predicted MW of the tagged protein variants are indicated. **B.** Western blot analysis using α-Ty antibody to detect tagged CAP5.5V (left) and CAP5.5 (right) proteins in bloodstream form (BF) cells. The α-Ty blot shows expression of both mT2A::3xTy and 3xTy – tagged variants and T2A-mediated release of 3xTy-tagged puromycin (3xTy::PAC) protein. Parental cells (Par) serve as negative control. **C.** Localisation of CAP5.5V::mT2A (CAP5.5V::mNG) and CAP5.5::mT2A (CAP5.5::mNG) in BF stages visualised by live-cell epifluorescence microscopy (green), co-stained with Hoechst 33342 (magenta; DNA). CAP5.5 is below detection limit in BF stage cells. Arrowhead indicates no staining of the flagellum. Scale bar = 5 μM. **D.** Flow cytometry analysis of mNG fluorescence in CAP5.5V and CAP5.5-tagged cells before and after differentiation from BF to PF life cycle stages. **E.** Quantification of flow cytometry results in D, showing median fluorescence intensity for mNG in BF and PF stages.

Live-cell microscopy confirmed the characteristic cytoskeletal localisation of CAP5.5V across the cell body (excluding the flagellum) in bloodstream cells (**Fig. 7B**), while CAP5.5 expression was not detectable above background, consistent with its known stage-specific expression. Anti-Ty immunoblots revealed stoichiometric expression of the tagged proteins: CAP5.5V::3xTy::T2A (104 kDa), CAP5.5V::mNG::3xTy::T2A (131 kDa), CAP5.5::3xTy::T2A (100 kDa), and CAP5.5::mNG::3xTy::T2A (127 kDa), as well as the cleaved 3xTy::PAC product (∼25 kDa), indicating efficient T2A processing (**Fig. 7C**). After differentiation to procyclic forms, flow cytometry showed the expected stage-specific regulation: CAP5.5V fluorescence decreased, while CAP5.5 signal increased (**Fig. 7D** and **7E**), corresponding to the previously reported 19-fold upregulation of CAP5.5V mRNA in bloodstream forms and 5-fold enrichment of CAP5.5 mRNA in procyclic forms (8). This differential expression of the two paralogs before and after differentiation validates the ability of the pRExT2A system to preserve complex stage-specific gene expression patterns mediated by endogenous UTRs.

## Discussion

Kinetoplastids such as *T. brucei* serve as powerful models for understanding fundamental biological processes, particularly the diversity and evolution of eukaryotic gene control (1, 3). The genetic toolbox for *T. brucei* is extensive, with existing tools for genome-wide conditional expression and localisation screens (24, 50, 51). However, efficiently tagging proteins in their native regulatory context, with positive selection for high throughput studies, remains challenging.

Our new CRISPR/Cas9-T2A system using the pRExT2A plasmids adds to this toolbox, enabling precise, efficient, scalable, stage-specific expression while retaining native mRNA processing signals, specifically native spliced leader addition sites (SLAS), native polyadenylation sites (PAS), native untranslated regions (UTRs), and any regulatory elements contained within these regions (1, 19). Importantly, the tagged proteins tested in this system retain correct localisation and differential expression in the tested life cycle stages. We demonstrate the versatility of this system by tagging proteins involved in key pathways, including GPI-PLC - which is critical for membrane lipid remodelling (52), CAP5.5V and CAP5.5, which are involved in cytoskeletal organisation (7, 8), and ESAG7, a subunit of the transferrin receptor responsible for iron uptake (53). The high efficiency allows for tagging up to 10 proteins per week in bloodstream form parasites, as the protocols for generating repair templates are well-established (25). A beginner’s guide for using this toolkit is outlined in **Fig. 8**.

**Fig. 8:**
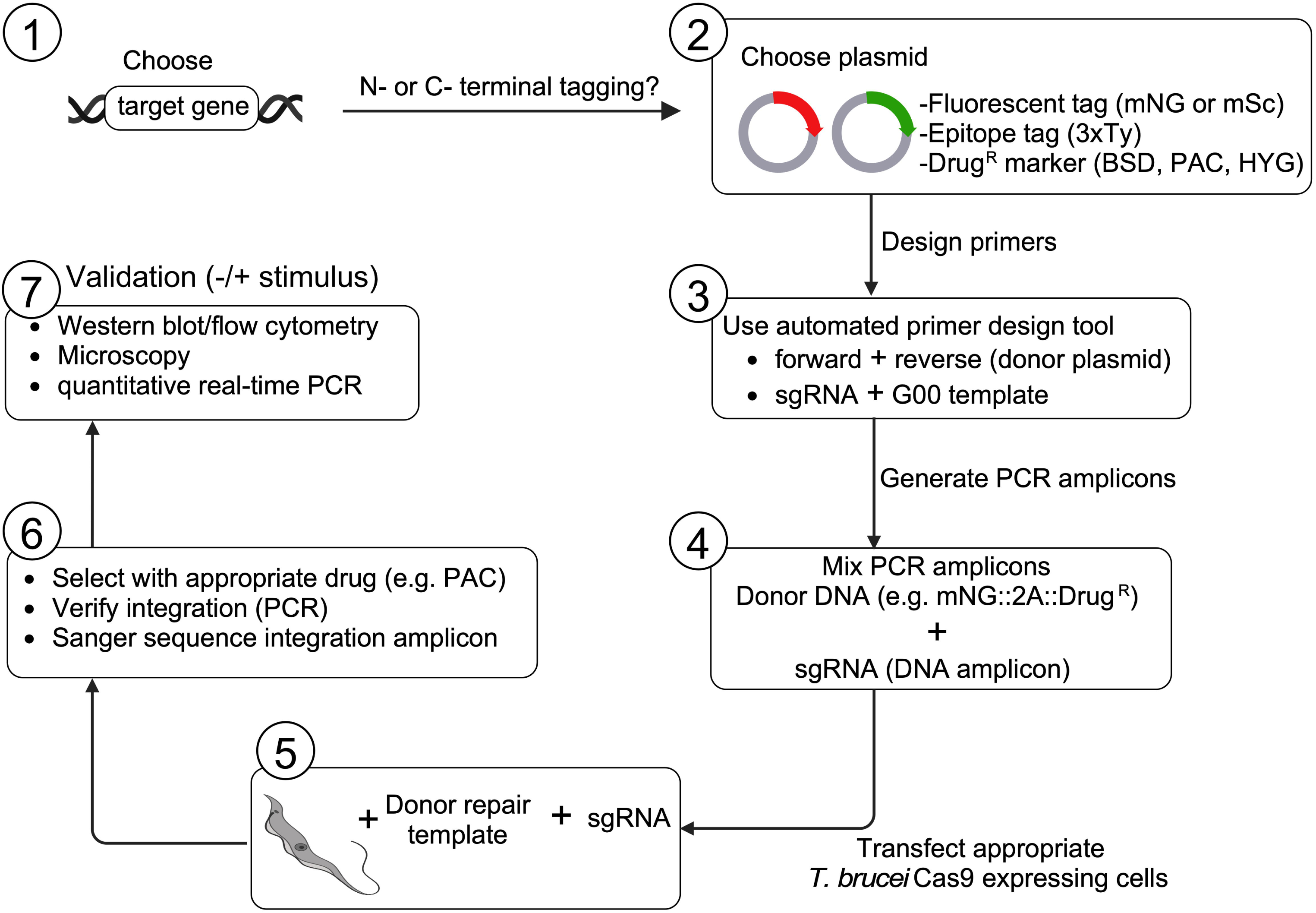
Beginner’s guide to using the pRExT2A tagging system: Start by selecting your target gene and deciding whether to tag it at the N- or C-terminus, or both. Next, choose the appropriate plasmid containing the desired fluorescent or epitope tag, along with a compatible drug-resistance marker (e.g., Blasticidin or Puromycin). Design primers using the automated tool to generate PCR amplicons for both the donor repair template and the sgRNA. Once the donor template and sgRNAs are ready, mix and transfect into *T. brucei* cells expressing Cas9. After transfection, apply drug selection to isolate successfully tagged cells. Validate the transgenic cells by performing PCR and Sanger sequencing to confirm correct integration. Finally, verify protein expression and localisation using Western blot, flow cytometry, or microscopy. Created with BioRender.com.

A key innovation of our system is the dual functionality of the drug resistance marker as both a selection tool and an expression control (**Fig. 5**). This feature allows users to distinguish between transcript abundance/translation efficiency and post-translational events by comparing the levels of the resistance marker to the tagged protein of interest. This approach has proven valuable in related kinetoplastids, such as *L. mexicana*, where T2A-based tagging showed that life cycle-dependent expression of LmxGT1 is controlled by changes in protein stability rather than mRNA levels (32). In contrast, in *L. donovani*, T2A applications showed that upregulation of LdNT3 is driven by increased transcript stability and translational efficiency, not changes in protein turnover (34).

Using this system in *T. brucei*, we can systematically study proteins involved in nutrient transport and their regulation across different stages of the life cycle. For example, this system can enable investigation into the multi-layered control of iron response genes by examining multiple proteins identified from transcriptomics and proteomics screens (10). This approach can help distinguish between transcript abundance, translation efficiency, and post-translational regulatory events (**Fig. 5D**), while characterising iron response elements (IREs) in a high-throughput format. Furthermore, combining our system with single-molecule RNA FISH enables simultaneous analysis of RNA processing dynamics, offering insights into gene regulation at all levels in a single experiment.

Another advantage of this system is its compatibility with the pPOT platform, enabling seamless integration into existing workflows and allowing for direct comparisons with protein expression levels where exogenous UTRs from the pPOT system serve as controls (25). For instance, we show that tagging ESAG7 by swapping UTRs diminished native regulation and function (**Fig. 4).** The modular architecture of the pRExT2A offers flexibility for easy swapping of tags, drug resistance markers, or regulatory elements without the need for extensive re-cloning. This adaptability facilitates rapid development of new constructs for diverse experimental applications.

A key strength of this system is its ability to track stage-specific protein expression before and after differentiation, as demonstrated for the dynamic regulation of cytoskeletal proteins CAP5.5V, and CAP5.5. By tagging multiple components with different fluorophores, we can visualise complex assembly and disassembly in real-time during differentiation, revealing both the timing of protein interactions and factors affecting complex formation. For example, tagging components of mitochondrial respiratory complexes alongside iron-sulfur cluster assembly proteins could allow tracking of respiratory complex assembly relative to transferrin receptor (TfR) downregulation. Such experiments could reveal how iron is mobilised from storage proteins to newly synthesised mitochondrial complexes and how iron availability affects these metabolic transitions.

Our T2A system has several technical considerations. First, CRISPR-based integration leaves a small (20 bp) scar sequence that could theoretically affect gene expression, though we observed no impacts during validation experiments. To address this potential limitation, we provide pre-designed primers for scarless integration, although these remain to be tested. Second, genome-wide analysis of PAM sequence requirements in 9,666 *T. brucei* genes shows that 87.6% have suitable N-terminal PAM sites and 79.3% have C-terminal sites. Given the GC content of *T. brucei* TREU927 (45.47%) and Lister427-2018 (43.67%), NGG PAM sites theoretically occur every 8-10 bp, providing multiple targeting options near gene termini (54, 55). Switching termini usually resolves targeting challenges when optimal sites are unavailable. Third, the system predominantly tags one gene copy, which can be challenging when targeting genes within multigene arrays that share identical UTRs, such as *Hsp70*. For such cases, specific targeting can be achieved by manually designing primers to regions that are not identical within the UTRs.

Fourth, when tagging developmentally regulated genes, users should evaluate available mRNA and protein data to guide experimental design. Our data with CAP5.5 demonstrates successful tagging despite rapid protein turnover in bloodstream forms (t_1/2_ = 2.3 h versus median t_1/2_ = 5.6 h) (56). The relatively stable mRNA half-life (33 min vs. median BSF mRNA t_1/2_ of 13 min) and detectable ribosome association enable sufficient expression for drug selection (57, 58). However, genes with highly unstable mRNAs in the chosen life cycle stage might be challenging to tag, as drug resistance and protein expression are controlled by the same regulatory elements. We recommend consulting published mRNA stability and ribosome-profiling data when designing T2A tagging strategies for stage-specific genes (2, 57, 58), keeping in mind that some genes may be successfully tagged despite low predicted expression.

For protein detection, the shared 3xTy epitope tag between the drug resistance marker and target protein creates specific constraints. While Western blotting can distinguish these proteins by size, immunofluorescence cannot separate their signals. We recommend either live imaging or using anti-mNeonGreen antibodies for localisation studies and choosing drug resistance markers of a different molecular weight to your target protein. Additionally, the limited dynamic range of mNeonGreen may not detect subtle expression changes, such as responses to nutrient starvation. To overcome this, we are developing luciferase-based vectors that offer enhanced sensitivity for monitoring dynamic responses to environmental signals.

Our T2A system has the potential to offer new insights into the complex biology of these parasites and contribute to our broader understanding of eukaryotic gene regulation, particularly in systems where post-transcriptional control plays a dominant role. By preserving native regulatory contexts, this approach could accelerate our understanding of molecular mechanisms underlying parasite adaptation and survival.

## Materials and methods

### *T. brucei* cell culture and generation of constitutive Cas9 expressing cells

A tetracycline inducible single-marker bloodstream form Lister 427 strain *T. brucei brucei* was used for all experiments (59). The cells were transfected with a humanised *Streptococcus pyogenes* Cas9 plasmid (pTB011) that integrates upstream of the first β-tubulin gene in the tubulin array and is under Pol II transcription (37). Cas9 expression was confirmed by Western blot using anti-Cas9 antibody (Abcam). The cells were cultured at 37°C, 5% CO_2_ in HMI-9 medium supplemented with 20% heat inactivated Foetal Bovine Serum, 100 U/ml penicillin and 100 μg/ml streptomycin (Gibco). Cell lines were propagated with the appropriate selection drugs where necessary: 2.5 μg/ml G418 (parental) and 10 μg/ml Blasticidin for Cas-expressing cells and 0.2 µg/ml Puromycin for reporter or epitope-tagged cell lines.

### Construction of T2A peptide-based plasmid templates for endogenous tagging

The “donor vectors” for T2A peptide-based tagging were originally designed as “SfiI cassettes” to fit into our multi-fragment assembly method for generating targeting constructs for integration via homologous recombination (60). The PAC::T2A::mNG and mNG::T2A::PAC tagging cassettes were synthesised in full by GenScript. The PAC and mNeon genes were encoded with a *Leishmania donovani* codon bias (i.e., predominantly G or C in the wobble position), and a nine amino acid flexible linker was included upstream of the T2A peptide sequence to facilitate efficient co-translational cleavage (38) The mScarlet (61) gene was PCR amplified from pCytERM_mScarlet_N1 (Addgene_85066) and inserted into N-terminal and C-terminal “Empty T2A::Drug^R^ cassettes as described in the **Fig. S1**. All pRExT2A plasmids were synthesised by Genscript using sequences from the pPOT series which have been optimised for *T. brucei* (25).

### sgRNA and repair template design and assembly

All oligonucleotides for generation of donor DNA repair templates and for analyses of edited genomic loci were synthesised by Life Technologies (Invitrogen, UK) and presented below or in the supplementary material. sgRNA primers targeting specific genomic loci were selected by choosing a 20 bp sequence adjacent to a Protospacer Adjacent Motif (PAM, 5’-NGG) site that is either close to the start codon (for 3’ targeting) or close to the STOP codon (for 5’ targeting). SgRNA primers were cross-checked for any off-target binding. PCR amplification of all sgRNAs followed the protocol published Beneke et al using 2 μM of primer G00 (sgRNA scaffold) as template and a gene-specific sgRNA primer (37). The resulting sgRNA PCR product and matching PCR-amplified repair template were pooled and transfected into cells that express T7 polymerase for T7 RNAP-driven transcription (59).

The repair template primers were designed to amplify the fluorescent TAG::T2A::drug^R^ with 25 - 40 bp homology arms targeting the gene of interest and the adjacent UTR (Fig 1). The homology arms were selected such that full length endogenous 5’ and 3’ UTR sequences remained intact following repair. The STOP and START codons were omitted for either C-terminus or N-terminus tagging, respectively. To disrupt the PAM sequence or sgRNA binding regions within the repair templates synonymous mutations or an *Xba*I site was introduced in the repair template primer, highlighted in bold in the primer sequences (26). For PCR amplification of repair templates, 30 ng of plasmid was used with gene-specific forward and reverse primers (**Table 2**).

**Table 2:**
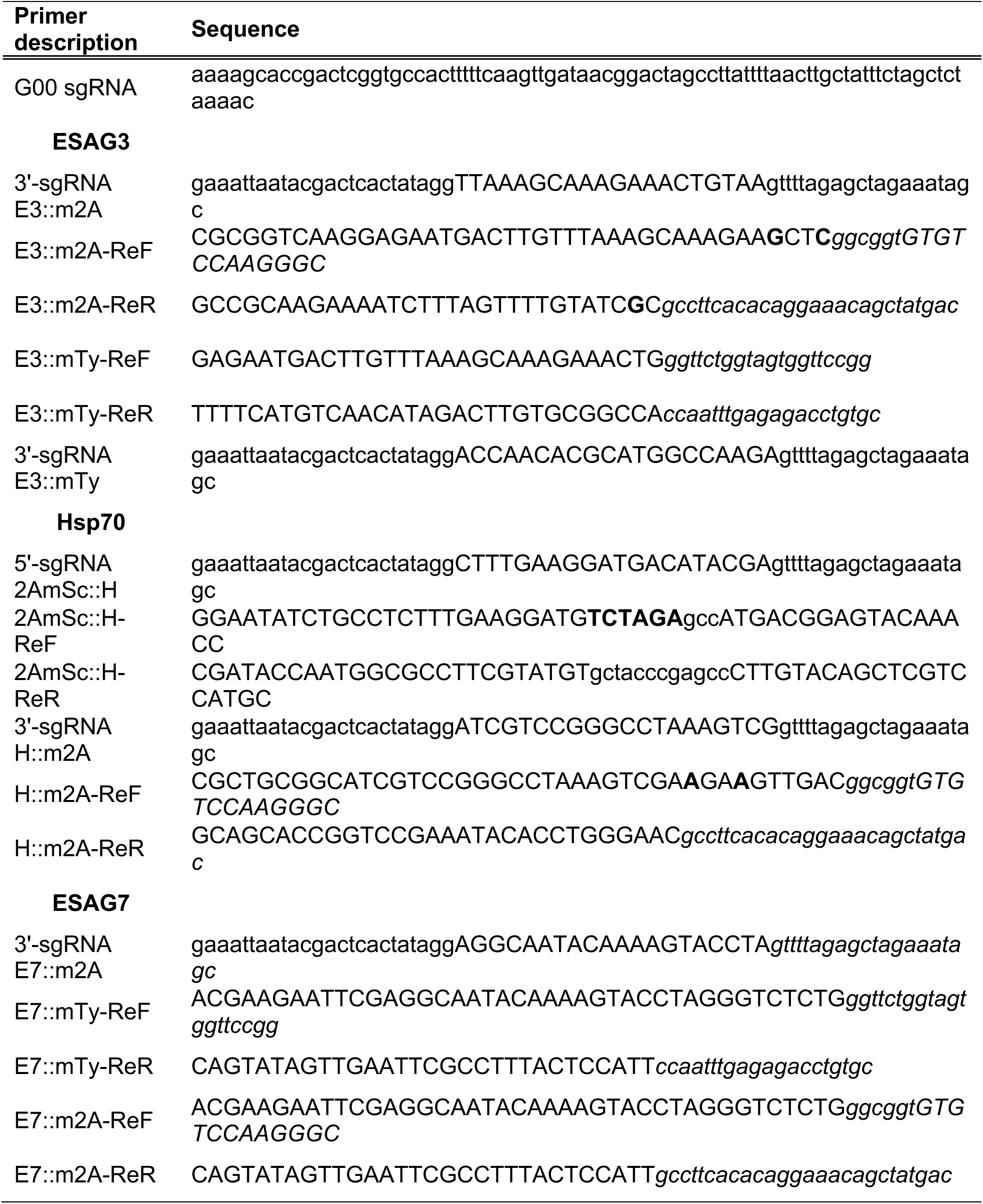
Primer sequences used for PCR amplification of sgRNA and DNA repair templates. ReF; forward repair and ReR; reverse repair. Primer binding sites corresponding to regions in the plasmid are shown in italics. Bold indicate synonymous mutations to prevent further Cas9 edits.

### PCR of repair templates and analyses of genomic integrations

All PCR assays for amplification of repair templates were performed in a 50 μl reaction volume using Q5 High-Fidelity 2X MasterMix (New England Biolabs NEB Cat #M0515) according to manufacturer’s instructions. The reagent components were assembled on ice and transferred to a preheated thermal cycler at 98.0°C. The annealing temperature for each primer pair was calculated using the NEB Tm calculator online tool. To validate correct integration of tags at the target loci, DNA was extracted from wild type and transfected cells using a DNeasy blood and tissue Kit (Qiagen). All PCR assays for genomic DNA analysis of edited loci were performed in a 25 μl reaction volume using Hot Start Taq 2X Master Mix (NEB, Cat #M0496). Five microlitres of all PCR reactions were resolved on a 1% agarose gel to verify the presence of correct size PCR products. For high-throughput applications, we recommend well-established workflows published by Beneke et al 2017 (25).

### Transfection of pooled PCR-amplified sgRNA and donor repair templates

1-3 x 10^7^ Lister 427 bloodstream cells were harvested and resuspended in 1x TbBSF Roditi buffer premixed with 50 µL of pooled PCR reactions (25 μl repair template fragment + 25 μl sgRNA; ∼25 μg total DNA each) without purification. The cell suspension in a final volume of 200 μl was electroporated using an Amaxa Nucleofector IIb (Lonza), program X-001. Cells were recovered for 6h at 37 ^0^C/5% CO_2_ before selection with appropriate antibiotics and distributed in 24 well plates (2 ml in each well). Positive clones were selected after 5 – 7 days and analysed by flow cytometry and Western blots. Correct integration of repair templates at target loci were confirmed by locus-specific PCR and genotyped by performing whole genome sequencing at BGI Genomics (China). Locus-specific PCR primers are detailed in **Table S1**.

### Western blotting and antibodies

Total protein was extracted from wild type and transgenic cells, washed in 1x Phosphate buffered saline, lysed in 1x SDS loading buffer, separated by SDS PAGE, transferred to PVDF membranes and immunoblotted with appropriate antibodies for 1 hr at room temperature: anti-ESAG3 (rabbit, 1:2500), anti-TfR (rabbit, 1:2500), anti-Hsp70 (rabbit, 1:5000) (62), anti-mNG (Mouse monoclonal antibody [32F6], ChromoTek 1:1000), anti-RFP (Mouse monoclonal antibody [6G6], ChromoTek 1:1000) and anti-2A (monoclonal clone 3H4, Merck, 1:1000). The membranes were blocked in 5% milk, washed 3x in 1xPSBT after antibody incubation and incubated for 1 hr at room temperature with the appropriate horseradish peroxidase (HRP)-conjugated secondary antibodies (anti-mouse IgG, 1:5000; and anti-rabbit IgG, 1:5000 dilution), developed after 1 min incubation with Super Signal ECL reagent (Thermo Fisher Scientific), and imaged using a ChemiDoc Gel Imaging MP System (Bio-Rad).

### Quantitative real time PCR (qRT-PCR)

All qRT-PCR analyses used ∼1–5 x 10^7^ cells total, and RNA was extracted from cell pellets using Qiagen RNeasy Extraction Kit. qRT-PCR from total RNA was performed using Luna Universal One-step kit (NEB) per manufacturer’s instructions. All reactions were performed in technical duplicates and specificity of each reaction was checked by ensuring single peaks from melt curves. Fold change was calculated by comparative C_T_ method using the Applied Biosciences software. Relative transcript levels were compared to endogenous control, *ZFP3* (Tb927.3.720, nts 241-301) relative to untreated controls. Primer sequences are available in supplementary material.

### Heat shock and iron starvation inductions

Iron starvation was induced for 5 hr at cell densities 5×10^5^ cells/ml with 25 mM Deferoxamine (Sigma Aldrich) in HMI-9 medium. For heat shock, Hsp70 tagged cells were maintained at 37 °C or transferred to 42 °C for one hour in HMI-9 medium at same cell densities £ 5×10^5^ cells/ml. In both cases, cells were harvested and analysed by qRT-PCR, flow cytometry or Western blots.

### Immunofluorescence microscopy and DAPI staining

Immunofluorescence assays were performed as previously described in (63). αBiP (Rabbit) antibody dilution was 1:2000, anti-mNG (Mouse monoclonal antibody [32F6], ChromoTek 1:100). Slides were washed with 1x PBS then incubated with secondary antibody for 1 hr at room-temperature: Alexa-488 conjugated a-mouse (1:500) and Alexa-594 conjugated a-rabbit (1:500) (Molecular Probes, USA). Slides were washed in 1x PBS and mounted in VectaShield mounting medium (Vector Laboratories) containing 4′,6-diamidino-2-phenylindole (DAPI), sealed with coverslips and imaged on a Zeiss AxioImager M2 with an ORCA Flash 4 camera.

### Flow cytometry

Live cells were harvested, washed in 1x PBSG, resuspended in 1 mL 1x PBSG, and analysed with an LSR Fortessa cell analyser using BD FACS Diva software (BD Biosciences). For uptake of Alexa-647 transferrin (Tf-647), the cells were pre-incubated with serum-free HMI9 supplemented with 0.5 μg/mL BSA at 37 ^0^C for 10 minutes. Tf-647 was added at 50 μg/mL and incubated for 30 minutes. The cells were washed twice in 1x PBSG and analysed by flow cytometry. Post-acquisition analysis of flow cytometry data was performed with FlowJo v10 software (BD Biosciences).

### Whole Genome Sequencing

Genomic DNA was extracted from wild type and Cas9-edited cells using the DNeasy Blood and Tissue Kit (Qiagen). Paired-end whole genome sequencing was performed by BGI Genomics (China) on a DNBseq platform with 100bp read length (PE100) for five tagged cell lines - ESAG3::mNG::T2A, Hsp70::mNG::T2A, ESAG7::mNG::T2A, ESAG3::mNG::Ty, and ESAG7::mNG::Ty, - and wild-type (untagged). Libraries were prepared using a BGI genomics custom short-insert DNA library construction protocol including DNA fragmentation, size selection, end repair and ’A’ tailing, adapter ligation, and PCR amplification. Raw reads were filtered using BGI genomics custom pipeline SOAPnuke (64) to remove adapter sequences, contamination and low-quality reads (parameters: -n 0.001 -l 10 -q 0.4 --adaMR 0.25 --ada_trim --minReadLen 100), yielding approximately 6.3-6.4 million clean reads per sample (1.26-1.29 GB data). Quality control of raw reads was evaluated and showed consistently high quality across all samples (97.99-98.16% bases ≥Q20 and 43.48-43.73% GC content). Clean reads were mapped to the *T. brucei* Lister427-2018 reference genome using Rsubread. For analysing the repetitive Hsp70 array, we employed systematic stringency controls in our mapping approach: from very high stringency (unique mapping only, no mismatches allowed, no alternate mapping allowed) to low stringency (allowing reads to map to multiple locations with up to 3 mismatches). Coverage statistics were calculated using GenomicAlignments in R.

### Development of an automated primer design tool

To facilitate use of this tool, we developed code for automatic design of primers. This was implemented as a Python module, using sequence and metadata retrieval from TriTrypDBv68 (65) to allow design of primers from a gene ID or list of gene IDs as found in TriTrypDBv68. For ease of use, we also wrote an iPython/Jupyter Notebook that runs in Google Collaboratory. Anyone can visit this notebook (available at: https://github.com/zephyris/scarlesstagging) and design primers without any need for coding ability or installing software on a computer.

The automated primer design attempts to design primers which make absolutely no change to the 5′ (for N terminal tagging) or 3′ (for C terminal tagging) UTRs. Without a CRISPR-mediated double strand break, this could be done by providing homology arms for the homologous repair construct which are, for N terminal tagging, the end of the 5′ UTR and start of the ORF or, for C terminal tagging, the end of the ORF (without the stop codon) and the start of the 3′ UTR. For CRISPR-assisted homologous recombination, a double strand cut site defined by a protospacer adjacent motif (PAM) site of GG must be selected near the start (for N) or end (for C) of the ORF. However, any such site would also cut the repair construct. Therefore, we search for protospacer adjacent motif (PAM) sites within the ORF which can be modified to remove the PAM without changing the encoded amino acid. This search range is defined by the number of bases typically available for primer synthesis (100 bases) minus the length of the template plasmid annealing site (template dependent, around 18 bases) and the length of homology (25 bases), searching within the start or end of the ORF for N or C terminal tagging respectively. The PAM site closest to the start (for N terminal) or end (for C) of the ORF is selected. The final primer sequences for the generation of the sgDNA are then constructed for the automatically selected PAM site, and the forward and reverse primer sequences for the homologous repair construct constructed for the user-selected template plasmid and automatically selected PAM site. The reverse (for N terminal) and forward (for C) primers contain a single point mutation in the ORF which removes the PAM site and so prevents CRISPR-mediated cutting.

## Data Availability

The whole genome sequencing data generated in this study have been deposited in NCBI Sequence Read Archive under BioProject accession PRJNA1202750. Raw reads for all tagged cell lines (ESAG3::mT2A, ESAG3::mTy, ESAG7::mT2A, ESAG7::mTy, HSP70::mT2A) and parental controls are included. These data support the integration analysis presented in Figure S3.

All pRExT2A plasmid templates for N- and C-terminal tagging described in this study have been deposited at Addgene (https://www.addgene.org/browse/article/28252442/).

## Code availability

The automated primer design tool for our tagging system is freely available at https://github.com/zephyris/scarlesstagging, along with pre-designed primers for *T. brucei* (TREU927, Lister 427)) and selected species, available at: https://github.com/zephyris/scarlesstagging/tree/main/examples/py/tritrypdb-genomes-v68.

## Acknowledgements

pCytERM_mScarlet_N1 was a gift from Dorus Gadella (Addgene plasmid # 85066; http://n2t.net/addgene:85066 ; RRID:Addgene_85066). The authors are grateful to Professor Jay Bangs (University at Buffalo, USA) for generous anti-TfR and anti-Hsp70 antibodies, and Professor Keith Gull, FRS for his insightful discussions and advice on gene choice for developmental regulation and experimental design.

