## Supporting information for "A high-throughput protein tagging toolkit that retains endogenous UTRs for studying gene regulation in Kinetoplastids"

**Running title:** An endogenous tagging toolkit with UTR retention in Kinetoplastids

**Key words:** Kinetoplastids, gene regulation, CRISPR/Cas9, molecular tools, T2A peptide, endogenous tagging, untranslated regions (UTRs), differentiation, *Trypanosoma brucei*

**Open Biology DOI: 10.1098/rsob.20240334**

**Table of contents:**

1. Supplementary Figures S1-S4
2. Supplementary Tables S1

Supplementary Figures

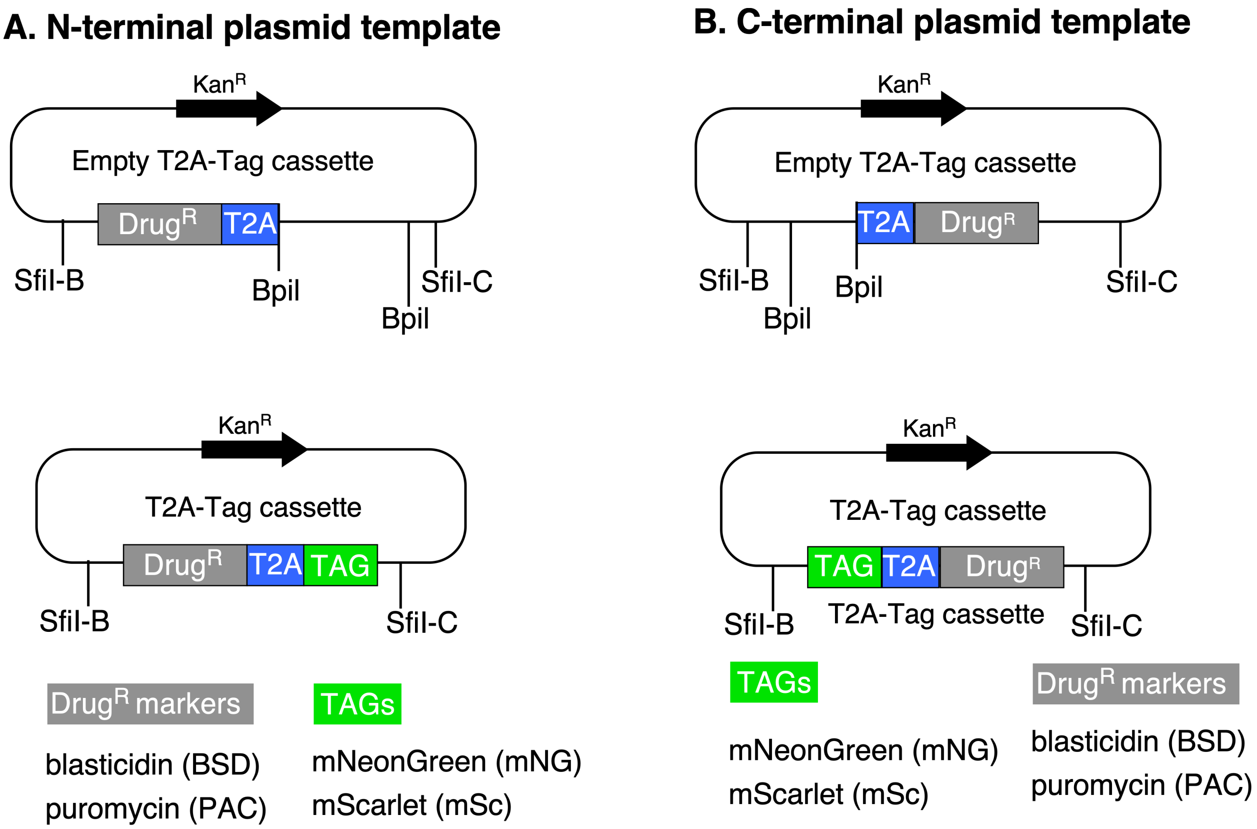


**Figure S1:** "**Empty T2A-drug cassettes**" consisting of the drug resistance gene (Drug$^R$) and N-terminal (**A**) or C- terminal (**B**) T2A peptide sequences, encoded with a *L. donovani* codon bias, were synthesised by Genscript. Two BpiI Type IIs restriction sites (removed upon cutting) allow seamless insertion of any desired tag to generate "**T2A-tag cassette**" donor vectors. Any gene or tag can be inserted into these vectors using compatible Type IIs restriction sites (BsaI, BpiI, BsmBI, etc.). For example, adding BsaI restriction sites with appropriate overhangs to primers for amplifying an open reading frame will facilitate in-frame cloning into the BpiI cut Empty 2A-drug cassettes (BsaI site is capitalised; overhangs generated upon BsaI digestion are underlined). PAC and BSD versions of N- and C-terminal GFP, ddFKBP, and ltDHFR tags have also been constructed. All T2A-tagging cassettes will be made available from Addgene.

N-Insert_BsaI-F: gcGGTCTCa tccg
N-Insert_BsaI-R: gcGGTCTCa ccga
C-Insert_BsaI-F: gcGGTCTCa cggt
C-Insert_BsaI-R: gcGGTCTCa agcc

The following primers were used to generate the mScarlet-2A-PAC and PAC-2A-mScarlet tagging cassettes, respectively:
N_mScarlet_BsaI-F: gcGGTCTCa tccg gtgagcaagggcgaggca
N_mScarlet_BsaI-R: gcGGTCTCa ccga cttgtacagctcgtccatgc
C_mScarlet_BsaI-F: gcGGTCTCa cggt gtgagcaagggcgaggca
C_mScarlet_BsaI-R: gcGGTCTCa agcc cttgtacagctcgtccatgc

### A. Schematic

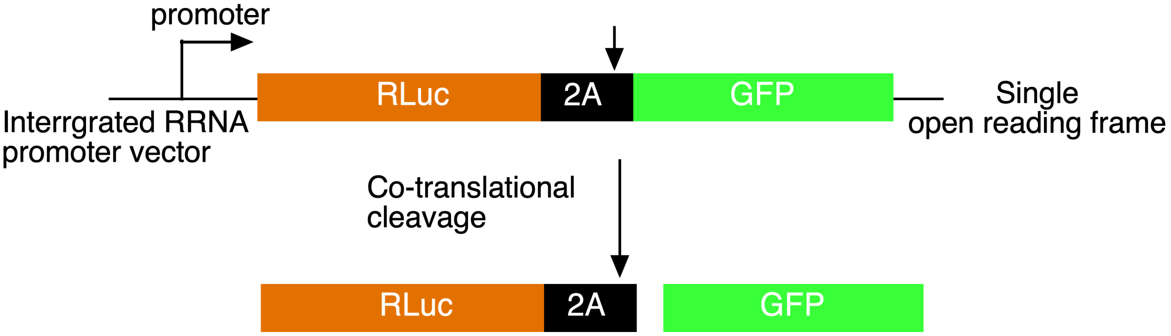


### B. Western blot

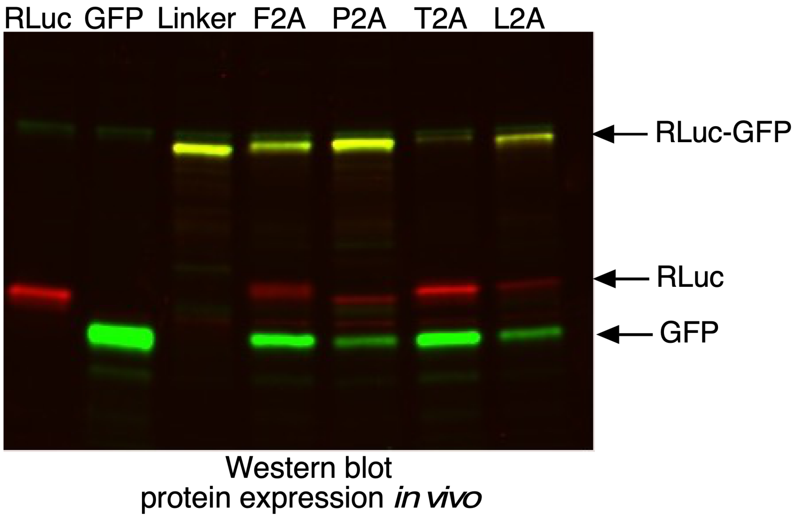


**Figure S2: The T2A peptide sequence promotes efficient co-translational cleavage in *L. donovani* promastigotes**

**A.** Schematic representation of a dual-reporter construct for co-expression of Renilla luciferase (RLuc) and green fluorescent protein (GFP), linked by a 2A self-cleaving peptide. The top panels illustrate construct designs where RLuc is fused to GFP either via a flexible amino acid linker or through different 2A peptide sequences, enabling co-translational separation of the two proteins. The 2A peptide sequences used were derived from: F2A (foot and mouth disease virus), P2A (*Porcine teschovirus*), T2A (*Thosea asigna* virus), and L2A (L1Tc retroposon from *T. cruzi*).

**B.** Western blot analysis of lysates from *L. donovani* promastigotes whole cell extracts expressing either RLuc, GFP, or an in-frame genetic fusion between RLuc and GFP separated by various 2A peptide sequences. Distinct bands corresponding to RLuc (red) and GFP (green) confirm successful efficient cleavage by the T2A peptide sequence, demonstrating efficient co-translational processing.

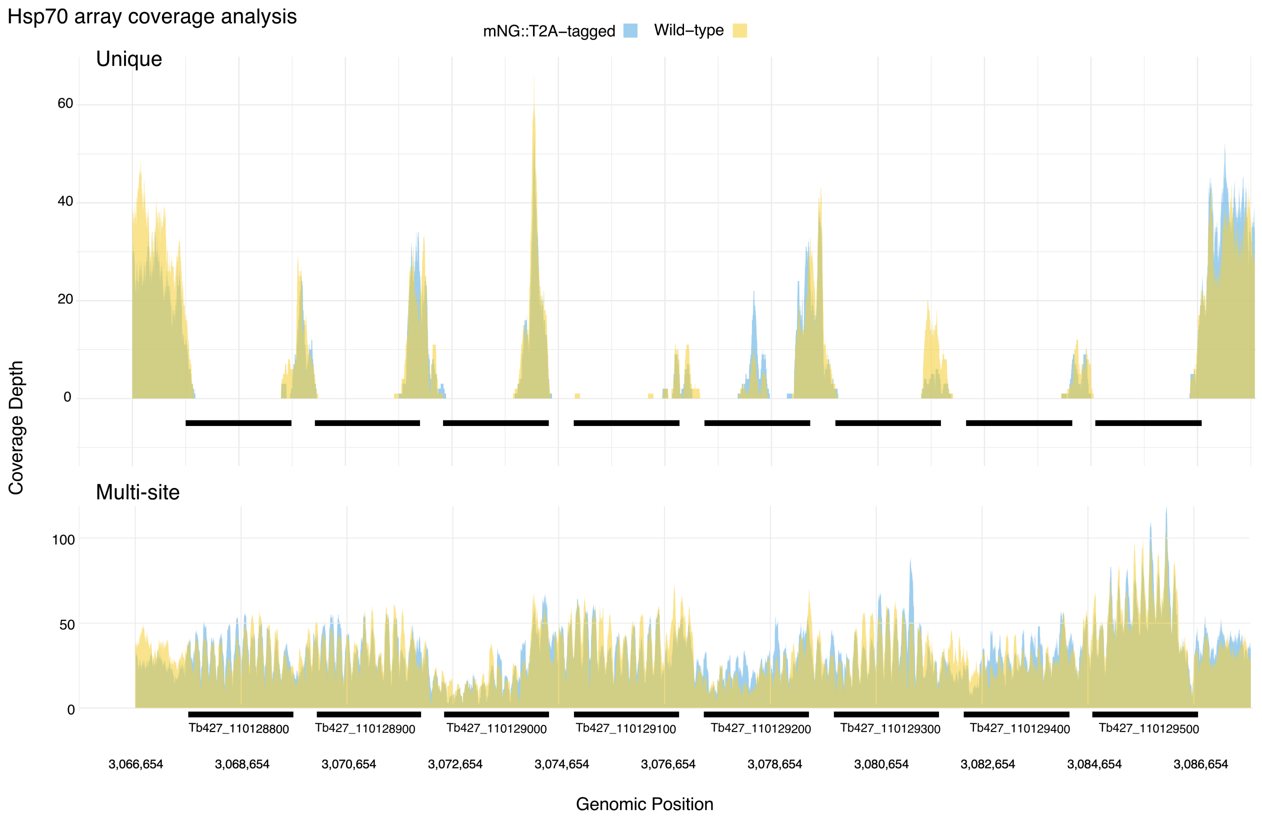


**Figure S3: Coverage analysis of the Hsp70 array using whole genome sequencing data mapped to the *T. brucei* Lister427-2018 reference genome (TriTrypDB-v68).**

Top panel: coverage when reads are restricted to unique mapping locations (no alternate mapping sites permitted, unique), revealing distinct regions within each gene copy. Bottom panel: coverage when reads are allowed to map to multiple sites, showing consistent coverage across all array copies (multi-site). Yellow tracks represent wild-type data, and blue tracks show Hsp70::mNG::T2A-tagged cell line data. Black bars indicate gene positions, gene IDs are shown at the bottom of each bar, genomic coordinates are shown below.

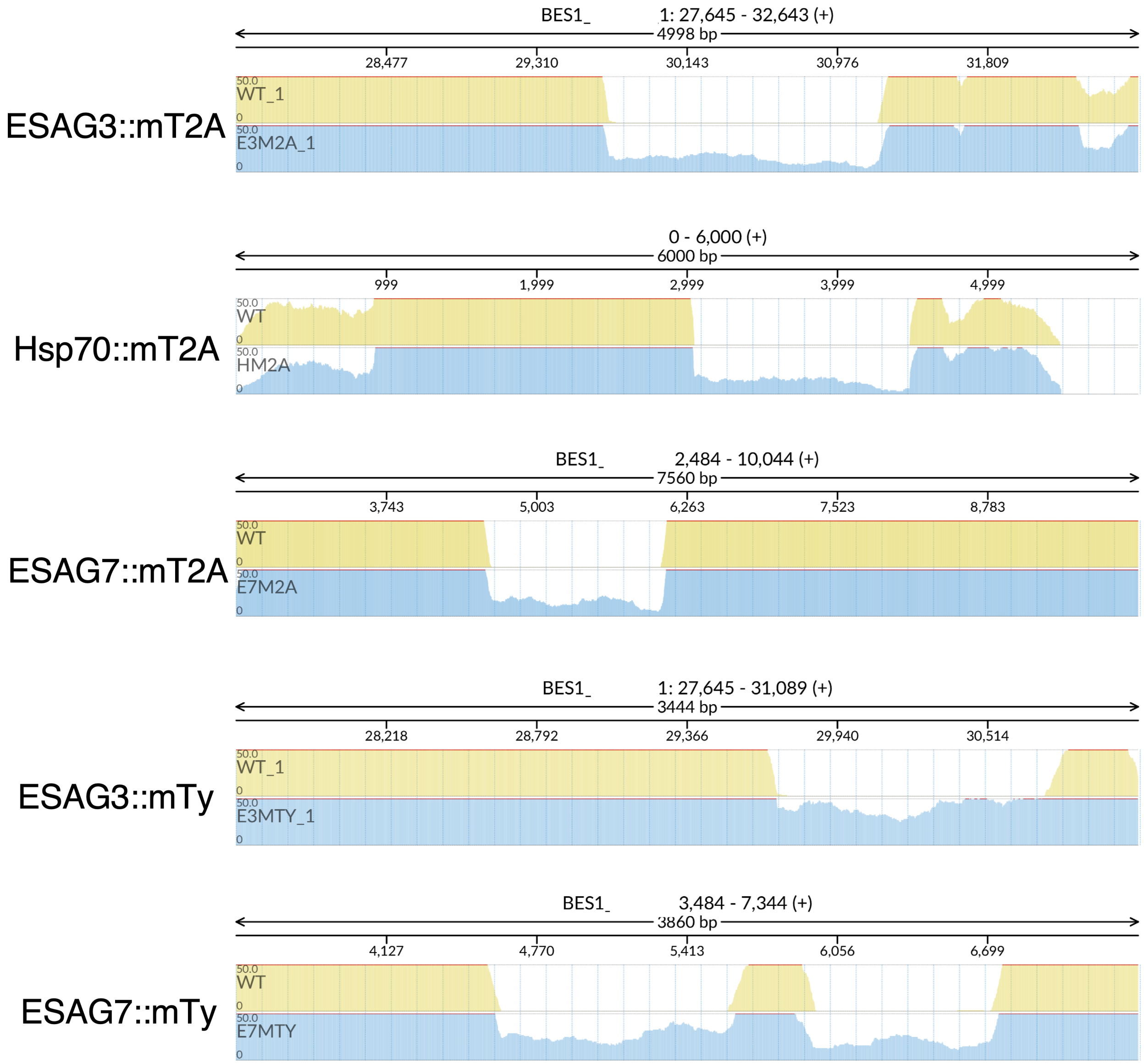


**Figure S4: Validation of fluorescent tagging cassette insertion at the target loci.** The plots show whole genome sequencing (WGS) read alignments for mNG-tagged genomic loci compared to wild type (WT) loci. The tagging cassettes were inserted at the following loci: ESAG3::mNG::T2A (ESAG3::mT2A), Hsp70::mNG::T2A (Hsp70::mT2A), ESAG7::mNG::T2A (ESAG7::mT2A), ESAG3::mNG::Ty (ESAG3::mTy), and ESAG7::mNG::Ty (ESAG7::mTy). Paired-end reads from each cell line were mapped to either the WT locus (yellow, upper track) or the manually reconstructed tagged locus (blue, lower track). Gaps in the WT locus correspond to the insertion of the mNG::T2A tagging cassette. In all cases, mNG::T2A integrated in-frame with the open reading frame (ORF) of the target gene and its endogenous untranslated region (UTR), with no mutations or frameshifts.

### Table S1: List of primers

| Primer ID | Primer Sequence | |
|---|---|---|
| CT932 | TGCCACCCACACCTTACCAATG | PAC-R for N-term tagging Hsp70 |
| CT754 | GTG ACA TTG CTG ACG ATT GAC | HSP70-qPCR-F |
| CT755 | AGA TCC TTG CCC TTG TTC TTG | HSP70-qPCR-R |
| CT49 | CAGGGGAAACGCAAAACTAA | TbZFP3 qPCR control primer |
| CT50 | TGTCACCCCAACTGCATTCT | TbZFP3 qPCR control primer |
| CT926 | ATACAACCACCGCCAGAAGGAG | Hsp70-ORFF For for C-term Hsp70 |
| CT927 | CGCATCTTTCCCGTAACGAGG | Hsp70-3UTR-R Rev for C-term tagging |
| CT928 | TGAAAACCGTAGCCAATGTGCG | mNEON-R for C-term tagging |
| CT929 | GCCTTCCTTGAAACTAGCGCA | PAC-F for C-term tagging Hsp70 |
| CT930 | ACGGTTACCCTGGTCATTTGCAAT | Hsp70-ORFR for N-term tagging |
| CT931 | CGTACTCCCATACTTGCACTCCTGC | Hsp70- 5'UTR-F for N-term tagging |
| CT932 | TGCCACCCACACCTTACCAATG | PAC-R for N-term tagging Hsp70 |
| CT933 | CACAACTGGCAACGGCAAGC | mNeon-F for N-term tagging Hsp70 |
| CT900 | GAGGAGTGGTGGTCAGAGCAGAG | Fwd ORF ESAG3 Tb427.BES40.16 |
| CT901 | GTAGCCAATGTGCGGGACAAGGATC | Rev primer for mNeonGreen |
| CT902 | TCACTGTGACCGCCGACGTG | Fwd primer for PAC. |
| CT903 | CAGCTTCTCGTGGTTTCTGCCTCAG | Rev primer for 3'UTR of ESAG3 |
| CT345 | GTGTGTGTGAGAGCTCTATTCC | Fwd primer for ESAG3 |
| CT1090 | TTCAGGGTGGCTGCATCCGATGCCTTTGAAATACCcG GTGATGAAACTGACAGTGATGAAggttctggtagtggttccgg | DownF-CAP5.5 |
| CT1091 | TTCTTGCATTTCTTTCTACTTATTGTATCCccaatttgagaga cctgtgc | DownR-CAP5.5 |
| CT1092 | gaaattaatacgactcactataggCCGATGCCTTTGAAATACgtttta gagctagaaatagc | DownSgRNA-CAP5.5 |
| CT1084 | TTTGCAGCTGATCGTATCGGTGCATTTGCCTTATCcGA TGATAGCGACACTGACggttctggtagtggttccgg | DownF-CAP5.5V |
| CT1085 | CTGTCTCTTATTTCCTTTCAGCCACCTCAAccaatttgagag acctgtgc | DownR-CAP5.5V |
| CT1086 | gaaattaatacgactcactataggTCGGTGCATTTGCCTTATgtttta gagctagaaatagc | DownSgRNA-CAP5.5V |
| CT1078 | GTTATGCGTTCATTTAAACACCAACCAAACCGCAAGaT CAggttctggtagtggttccgg | DownF-GPI-PLC |
| CT1079 | CAGCAGAGATTTGTTTTTTTTCCCACATTTccaatttgagag acctgtgc | DownR-GPI-PLC |
| CT1080 | gaaattaatacgactcactataggAACACCAACCAAACCGCAgtttta gagctagaaatagc | DownSgRNA-GPI-PLC |
